# Proteomic-based evaluation of nuclear transport of NLS-tagged trastuzumab-emtansine with enhanced cytotoxic potency

**DOI:** 10.1101/769588

**Authors:** Vincent Lacasse, Simon Beaudoin, Steve Jean, Jeffrey V. Leyton

## Abstract

Nuclear transport receptors (NTRs) are the only proteins able to transport large molecular weight payloads into the nucleus. A dominant area of molecular therapeutic research is the extension of the use of NTRs to target the nucleus for the development of pharmaceuticals or as tools for investigating fundamental biological questions. Although several examples of synthesized peptides harbouring nuclear localization signal (NLS) sequences conjugated to various payloads exist in the literature, the assumption has been that transport occurs by classical nuclear localization via the NTRs importin-α and importin-β. This assumption is relevant to nuclear-targeted therapeutics that aim for full potential clinical impact. In addition, fundamental research can benefit from unbiased approaches to investigate the role of NTRs. Herein, we report the construction of a novel NLS-modified agent composed of trastuzumab-emtansine (T-DM1) coupled to cell accumulator (Accum), a technology that enables monoclonal antibodies to escape endosome entrapment and accumulate conjugated payloads in the nucleus without abrogating affinity or specificity to target antigens. Accum harbours a classical NLS sequence from SV-40 large T-antigen. We demonstrate that routing T-DM1 to the nucleus successfully increased cytotoxic potency in the HER2-positive cell line SKBR3. More importantly, through the development of a novel bait-prey proteomic approach, we show that the non-classical NTR importin 7 and not importin-α/importin-β was required for the cytotoxicity effect. This result was validated by siRNA knock down. Our findings also indicate that by discovering an unanticipated NTR regulator of an NLS-modified agent, this study demonstrates the utility of combining an unbiased proteomic approach to probe NTR function in mammalian cell system and, is a foresight for future NLS-based development initiatives.

## INTRODUCTION

Signal peptides are an integral component of intracellular protein targeting. The application of nuclear localization signal (NLS)-tagging for nonviral transfer of oligonucleotides, proteins, and reporter molecules is widely investigated for the development of therapeutic medicines and as probes used for advancing fundamental biology. For example, nuclear targeting via NLS-tagging is improving efficient localization of DNA vaccines into the cell nucleus for subsequent expression of immunogenic peptides, which is an obstacle preventing clinical success (*1*). Nuclear targeting of theranostic radionuclides is being implemented in the clinic to apply short range-emitting radionuclides to impart high ionization densities at the site of decay, which maximizes DNA damage and could overcome much of the toxicity with traditional radiation approaches (*2*). Utilizing NLS-tagged recombinant reporter molecules as probes has facilitated discoveries on various aspects of normal and diseased cellular development (*3,4*). Hence, NLS-tagged agents are critical for future effective medicines as well as tools for the advancement of fundamental biology.

Nuclear transport receptors (NTRs), also known as karyopherins, mediate the transport of macromolecules across the nuclear pore complex (*5,6*). Because NTRs have this unique capability, they are essential regulators for many normal cellular functions. There are >20 NTRs in the human genome and many NTR-specific cargos have been identified creating multiple transport pathways. Recently, the role for NTRs has expanded beyond housekeeping functions in nuclear transport. Studies have demonstrated particular NTRs can be modified or increased in abundance in different species. The modification enables NTRs to localize to the plasma membrane and coordinate the scaling of intracellular structures, including the nucleus to cell size (*7,8*). In response to cellular stress, NTRs can accumulate in the nucleus to block nuclear protein import by directly binding nuclear components and can also directly exert gene expression (*9*). Transcription length variants of NTRs have also been shown to be preferentially expressed and have unique roles in cells at different developmental stages (*10*). In tumor cells, distinct NTRs have been shown to be upregulated (*11–13*). Thus, NTRs are now considered multifaceted whose regulatory duties are ever expanding in both healthy and diseased cells.

Although NTRs have been shown to transport specific cargoes (or cohorts of cargos), there is a general lack of consensus structures or NLS sequences for NTR binding. The classical NLS sequence PKKKRKV from the simian virus 40 (SV40) large T-antigen was the sequence identified and used to elucidate nuclear transport over three decades ago (*14–17*). Proteins that require entry into the nucleus contain an NLS that is recognized and bound by the adaptor protein importin-α. Importin-β then binds to importin-α and the complex is subsequently transported along the cytoskeleton by motor proteins and dock at the NPC. The cargo is released into the nucleus through the binding of Ran-GTP. However, most of nuclear transport is responsible by NTRs that can directly bind their cargos in the absence of importin-α (*18*). Thus, it is difficult to determine whether the cell may have particular reliance on any individual NLS and/or cargo.

Although NLS-tagged agents are established as important potential therapeutics and effective investigational tools, their design, and intent of use, is still largely based on the classical mechanism of nuclear transport. This report originated from our pondering whether NLS-tagged agents have been overly reliant on the classical import mechanics and on the NLS from SV40 large T-antigen. Major challenges blocking elucidation is that unique classes of NLS sequences of particular NTRs have not been identified and NTRs share similar molecular weights, isoelectric points, and have low sequence identity (*18*). In addition, cargos for the majority of NTRs have been identified through binary protein-protein interaction assays (*19–21*).

We set out to develop an approach to identify NTRs and test their uniqueness or redundancy to import an NLS-modified agent in live cells. We present the technology cell accumulator (Accum), which harnesses a two-step movement of endosome escape coupled to nuclear translocation (*22*). Accum is a natural composite compound of 1.8 kDa that is conjugated to surface lysines of monoclonal antibodies (mAbs). It is comprised of a single cholic acid linked to a short 13 amino acid peptide CGYGPKKKRKVGG containing the NLS from SV40 large T-antigen.

Accum addresses the current limitation with NLS-based agents. The barriers of NLS-tagged agents include target cell binding, internalization, endosomal escape, and nuclear localization. For therapeutic applications researchers are often limited by a significant trade-off – the major elements being high cell penetration/nuclear accumulation but loss of target cell specificity and vice versa. Accum-modified mAbs conjugated to radionuclides retain high target affinity and specificity yet can accumulate the delivered payload at increased intracellular levels relative to unmodified mAbs and mAbs modified with the NLS but no cholic acid (*22,23*). Following receptor-mediated internalization, Accum enables mAbs to escape endosomes by utilizing cholic acid to activate acid-sphingomyelinase and produce ceramide, which destabilizes endosomal membranes (*23*). This endosome escape feature results in increased efficiency in nuclear localization and accumulation of delivered payloads. As a result Accum-modified mAbs target tumors with superior tumor cell accumulation and specificity compared to non-Accum-modified counterparts (*24*).

Thus, we hypothesized whether Accum’s unique delivery mechanism could be applied as a therapeutic agent. We examined the effect of incubating human epidermal growth factor receptor 2 (HER2)-positive SKBR3 cells with the clinically approved biopharmaceutical trastuzumab-emtansine (T-DM1) modified with Accum (herein Accum-T-DM1). Our rationale was that T-DM1 effectiveness is reliant on successful tumor cell binding to HER2 on the cell surface, followed by internalization into the cell and encapsulation inside endosomes (*25*). Motor proteins then naturally transport endosomes to lysosomes for membrane fusion and transfer of the encapsulated contents (*26*). Lysosomal proteases digest the Tmab backbone of T-DM1 and liberate membrane impermeable DM1 metabolites that are presumably transported out of the lysosome by a lysosomal membrane transporter to diffuse throughout the cytosol, and encounter tubulin and inhibit microtubule polymerization (*27*). As a result, tumor cells undergo apoptosis. Although microtubules are present in the cytosol, Oroudjev et al., showed that breast cancer cells in the prometaphase/metaphase rather than interphase were more sensitive to antibody-DM1 conjugates (*28*). Thus, with Accum-mediated nuclear transport and the importance of the nucleus on DM1 cytotoxicity, Accum-T-DM1 could feasibly test our hypothesis. An accompanying objective was to evaluate Accum as a tool to probe nuclear transport regulation in live cells while maintaining target receptor specificity.

We constructed Accum-T-DM1 and ensured that it maintained biochemical and biophysical properties to develop as an effective NLS-modified agent. We conducted *in vitro* studies to evaluate nuclear localization and cytotoxicity with Accum-T-DM1 in the HER2-positive breast cancer cell line SKBR3. A pull down assay was developed involving treatment of living SKBR3 cells incubated with Accum-T-DM1, T-DM1, or Trastuzumab (Tmab) followed by mass spectrometry (MS). We organized our proteomic data into novel bait-prey models to extract proteins that specifically interact with Accum-T-DM1 with high certainty. We then conducted cytotoxicity assays in SKBR3 cells depleted for identified NTRs. Our findings showed that Accum-T-DM1 could be constructed as an NLS-modified agent for *in vitro* testing and that it increased cytotoxicity by several-fold relative to T-DM1 and was specific for HER2. Importantly, our findings showed that Accum-T-DM1 was not reliant on the classical NTR complex for enhanced cytotoxicity. Accum combined with our proteomic and biochemical methods combine as a potentially readily available approach for the future development of NLS-based agents for therapeutic applications. In addition, Accum-modified mAbs are potential tools to utilize for investigating fundamental biological questions.

## RESULTS

### Construction of Accum-T-DM1 conjugates and biochemical and biophysical characterization

Accum was conjugated to T-DM1 using the polyethylene glycol (PEG)ylated succinimidyl-4-(N-maleimidomethyl)cyclohexane-1-carboxylate (SM(PEG)_2_) crosslinker (Fig. 1A). The reduced SDS-profile of T-DM1 showed distinct bands at approximately 25 kDa and 50 kDa for the light chains (LC) and heavy chains (HC), respectively (Fig. 1B). These molecular weights were in agreement with the theoretical and experimental LC and HC masses of T-DM1 previously reported (*29*). T-DM1 reacted with increasing ratios of SM(PEG)_2_ followed by Accum conjugation gave rise to two major bands consistent with the presence of LC and HC relative to T-DM1 (Fig. 1B). Both LC and HC bands in the Accum-modified T-DM1 conjugates migrated consistent with increased molecular weight. In addition, as ratios of SM(PEG)_2_-to-T-DM1 increased the resultant Accum-T-DM1 conjugates exhibited slower migration on the SDS-PAGE gel. The resulting higher HC bands indicated more Accum attachments compared to LC, which is consistent as a majority of lysines in the human IgG1 antibody are located in the HC (*30*). The Accum-T-DM1 conjugates were >90% monomers when 25- and 50-fold excess SM(PEG)_2_ crosslinker ratios were utilized. The calculated number of Accum moieties per T-DM1 is listed in Figure 1B.

**Figure 1.**
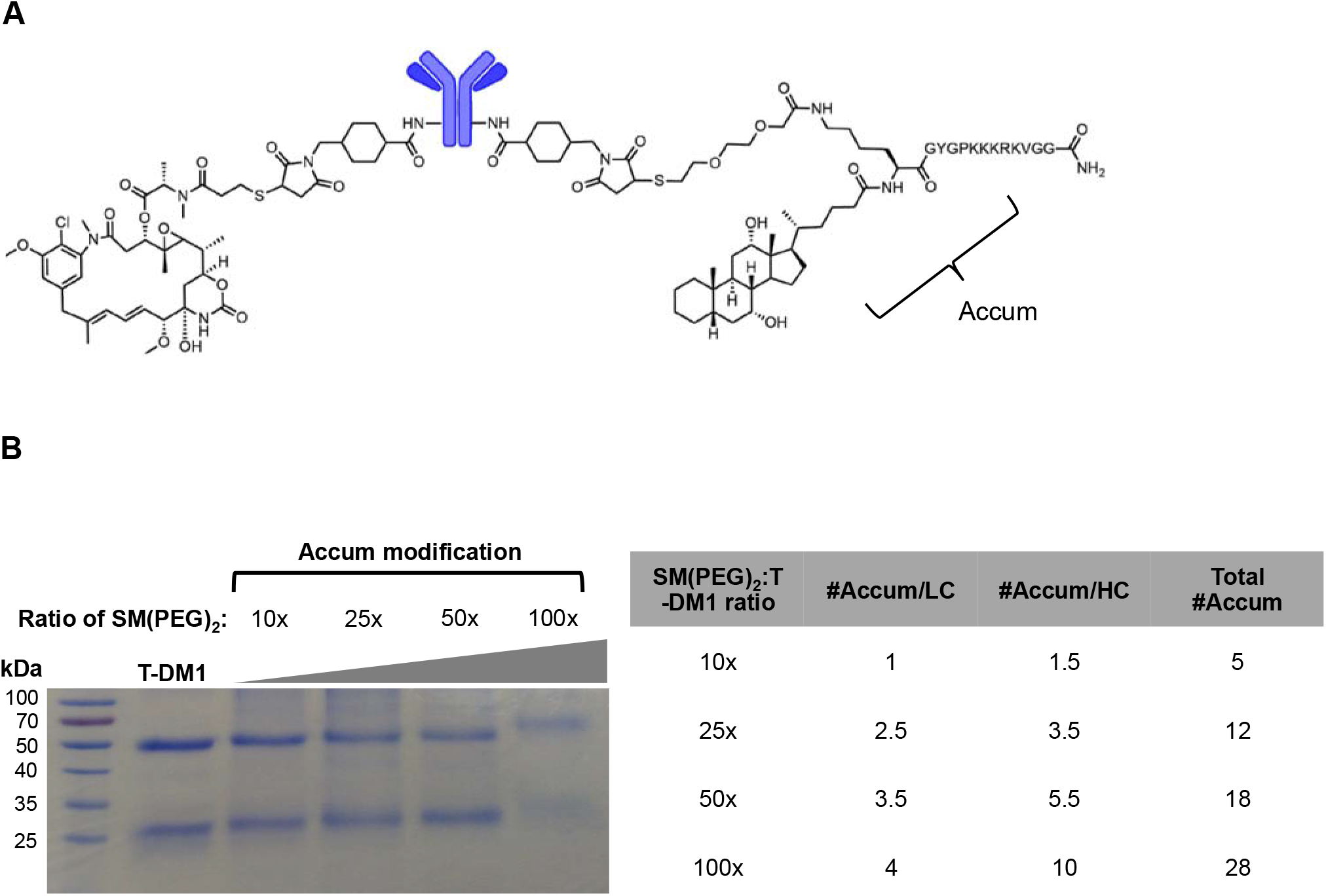
(A) Representative drawing of Accum-T-DM1 depicting lysine conjugation via SM(PEG)_2_. (B) Reducing SDS-PAGE of T-DM1 and Accum-modified T-DM1 at increasing SM(PEG)_2_-to-T-DM1 ratios. Table shows the calculated number of Accum moieties per light chain (LC) and heavy chain (HC).

To determine the Accum-T-DM1 stability profile, unfolding and aggregation were evaluated by label-free differential scanning fluorimetry (nanoDSF). Analyzing ADC stability with nanoDSF has proven effective for its ability to detect unfolding events of the mAb Fab, C_H_2, and C_H_3 domains (*31,32*). T-DM1 and T-DM1 modified with 12 and 18 Accum moieties, but not 28 moieties exhibited comparable starting levels in the initial ratio (350nm/330nm) (Fig. 2A). The thermal-induced unfolding transitions for T-DM1 C_H_2 and C_H_3 domains of the Fc region were 70.1 °C ± 0.2 °C and 79.5 °C ± 0.4 °C, respectively. In contrast, there was only a single unfolding transition for all Accum-T-DM1 conjugates. Accum_18_-T-DM1 and Accum_28_-T-DM1 ADCs had transitions at 70.4 °C ± 1.3 °C and 69.4 °C ± 0.3 °C, respectively. Accum_12_-T-DM1 had an unfolding transition at 73.5 °C ± 0.6 °C. These transition temperatures correspond to the unfolding of the CH2 domain. This study revealed an inverse correlation where increasing the number of SM(PEG)_2_-activated lysines with subsequent Accum conjugation reduced the temperature values for the CH2 unfolding transitions proportionally (Fig. 2B). This suggests that Accum modification has a greater impact on the CH2 domain compared to other domains of T-DM1. This is consistent with previous ADCs based on a human IgG conjugated via lysines to DM1 that showed the C_H_2 domain, relative to other domains, is the most likely to be destabilized due to a preference for DM1-lysine conjugations for C_H_2 (*32*). Based on theses results except for Accum_28_-T-DM1, the Accum-T-DM1 conjugates were soluble and had stabilities sufficient for functional evaluations in live cells.

**Figure 2.**
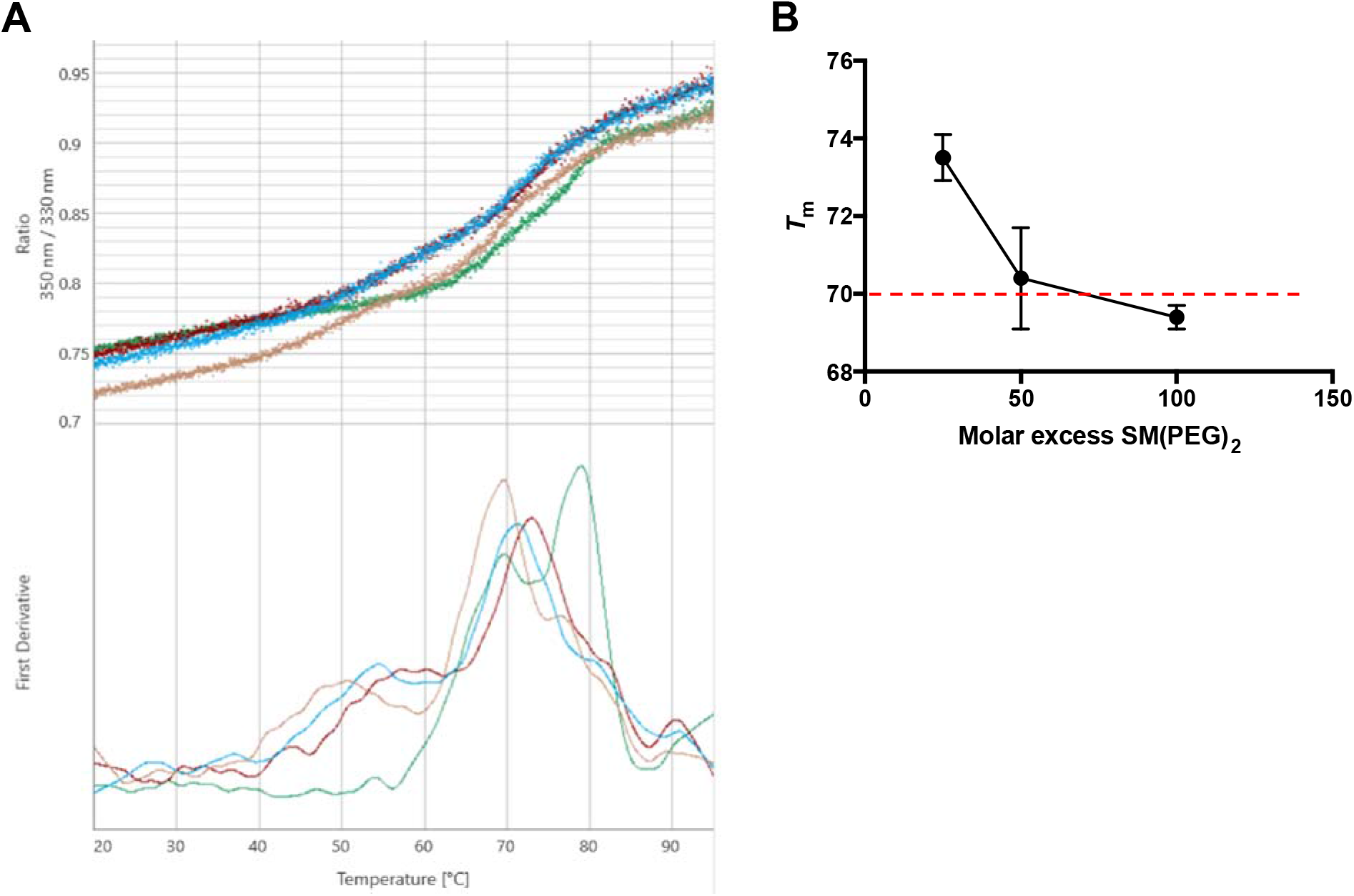
(A) Differential Scanning Fluorimetry thermograms of T-DM1 (green) and Accum-T-DM1 where T-DM1 was first reacted with 25- (red), 50- (blue), and 100-fold (orange) molar excess SM(PEG)_2_. Top, the 350 nm/330 nm fluorescence ratio versus temperature gradient. Bottom, the corresponding first derivatives are shown. (B) The *T*_m_ of the unfolding events plotted versus the molar excess of SM(PEG)_2_ used to react with T-DM1

### Accum-T-DM1 enhances cytotoxicity by localizing to the nucleus

Accum-modification enhanced T-DM1 cytotoxic potency. SKBR3 cells treated with T-DM1 resulted in an IC_50_ value of 0.22 nmol/L (R^2^=0.96). This result is amid the IC_50_ values of 0.047-0.12 nmol/L and 7 nmol/L reported by Lewis Phillips and Juntilla et al, respectively (*27,33*). In contrast, SKBR3 cells treated with Accum_6_-T-DM1, Accum_12_-T-DM1, and Accum_18_-T-DM1 resulted in an IC_50_ values of 0.024 nmol/L (R^2^=0.95), 0.020 nmol/L (R^2^=0.99), and 0.012 nmol/L (R^2^=0.99), respectively (Fig. 3A). To demonstrate specificity, HER2-negative MCF-7 breast cancer cells were treated with Accum-T-DM1 and T-DM1. There was no increased cytotoxicity against MCF-7 cells (Suppl. Fig. S1). These data indicate that an alternative delivery route for T-DM1 improved the cytotoxic effectiveness by 18-fold and did not abrogate specificity of T-DM1 for HER2.

**Figure 3.**
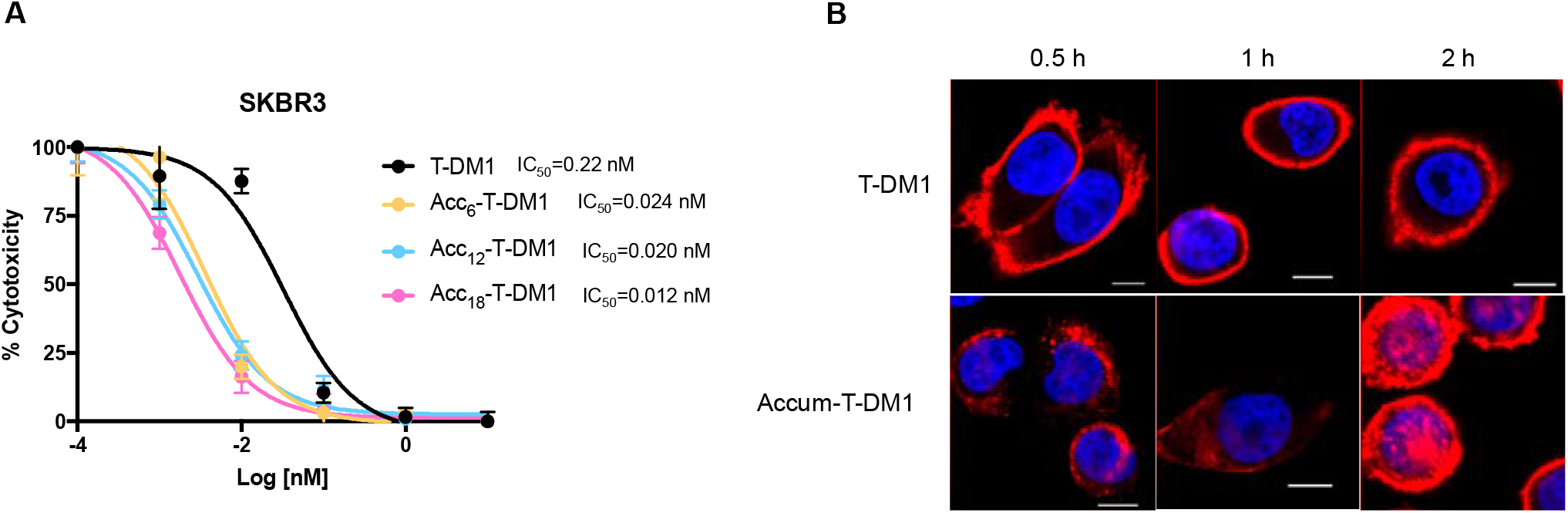
(A) Cytotoxicity profile of SKBR3 cells when incubated with increasing concentrations of T-DM1 and T-DM1 modified with increasing amounts of Accum (Acc). (B) Distribution of Tmab (red) in SKBR3 cells treated up to 2 h with Accum-T-DM1 or T-DM1 and then processed for immunofluorescence confocal microscopy. Nucleus in blue and scale bar is at 100 nm.

We aimed to evaluate nuclear localization of Accum-T-DM1 versus T-DM1 up to 2 h as internalization peaked at this time point and remained plateaued to the end of the 8 h evaluation time point (Suppl. Fig. S2). The confocal microscopy images showed that T-DM1 was exclusively localized at the plasma membrane (Fig. 3B). In contrast, confocal microscopy images showed that at 0.5 h Accum-T-DM1 was internalized and distributed in the cytoplasm shown as punctate foci typical of endosome entrapment. However, at 1 h Accum-T-DM1 distribution had increased diffusion in the cytoplasm. At 2 h, immunofluorescent signal pertaining to Accum-T-DM1 was clustered in the nucleus. There was also extensive signal from the cell surface in cells treated with Accum-T-DM1. This was most likely due to the expression level of SKBR3 as it is a standard for HER2 overexpressing breast cancers with 1 x 10^6^ receptors per cell or a 3+ on the clinical scoring system (*34*). In addition, a significant portion of Tmab (~75%) is known to remain at the cell surface for periods up to 18 h and removing HER2 on the surface of cells by traditional trypsin digestion or acid washing cannot be accomplished (*35,36*). Evaluations after past 2 h were not performed. However, studies in the HER2-positive cell line JIMT-1 show that the signal of Accum-T-DM1 is gradually decreased at later time points and is caused from degradation by nuclear residing serine and lysine proteases (unpublished). This experiment demonstrates that, indeed, Accum-modification of T-DM1 leads to nuclear localization. In addition, T-DM1 in the cytosol of SKBR3 cells could not be readily visualized throughout the evaluation suggests that the amount of T-DM1 that is able to enter the cell and be processed in the lysosome is rapid. In contrast, we could visualize Accum-T-DM1 in punctate vesicles at 0.5 h followed by diffusion in the cytosol at 1 h and nuclear localization at 2 h. This suggests a slower mode of intracellular drug delivery via routing to the nucleus versus the lysosome.

### Selecting the Accum-T-DM1 conjugate for proteomic experiments from treated live cells

As Accum-T-DM1 is composed of a mAb, payload, and Accum it is challenging to eliminate false positive protein interactions due to the structural complexity of this agent. An antibody affinity pull down with protein-G coated magnetic beads was chosen. The particular challenge with this type of pull down is that antibody-conjugates have potential binding sites to protein-G blocked due to lysines being occupied with Accum and DM1. Thus it was important to characterize the pull downs of lysates from various live SKBR3 cells treated with Accum-T-DM1 conjugates before progressing to the proteomic stage. As previously described, the maximum cellular uptake of the conjugates was at 2 h (Suppl. Fig. S2) and therefore was used as the incubation time point for the conjugates with SKBR3 cells. In order to use “physiologic” concentrations were explored various concentrations of Accum-T-DM1. A concentration of 7.5 mg/mL (~50 nmol/L) Accum-T-DM1 incubated with 15 x 10^6^ SKBR3 cells for 2 h was identified to provide sufficient antibody pull down. Figure 4 shows the pull down efficiency of Protein G-coated magnetic beads for the Accum-T-DM1 conjugates relative to Tmab and T-DM1. The amount of Tmab pulled down from treated SKBR3 cells was comparable to a 5 μg standard of Tmab added to Protein G-coated magnetic beads. Relative to the artificial Tmab standard, the amount of T-DM1 pulled down was noticeably reduced. The amount of all the Accum-T-DM1 conjugates pulled down was reduced relative to T-DM1. In fact, in cells treated with Accum_28_-T-DM1 there was no visual antibody present on the gel corresponding to the molecular masses of the HC and LC. Only Accum6-T-DM1 was pulled down at comparable, albeit reduced, levels relative to T-DM1. Based on these results, Accum_6_-T-DM1, for its ability to localize to the nucleus, enhanced cytotoxicity, and ability to be pulled down by Protein G-coated magnetic beads was the agent utilized for proteomic evaluation of nuclear transport.

**Figure 4.**
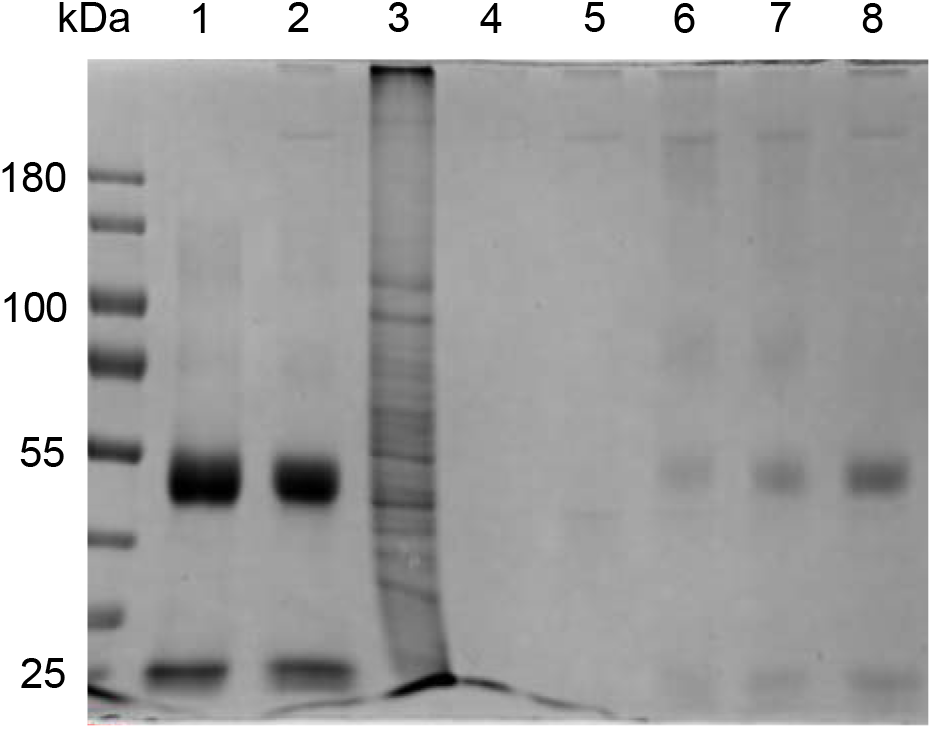
Representative Western blot from a reducing SDS-PAGE gel of SKBR3 cells treated with Tmab, Accum_28_-T-DM1, Accum_18_-T-DM1, Accum_6_-T-DM1, and T-DM1 shown in lanes 2, 5, 6, 7, and 8, respectively (n=5). Lane 1 is a pull down of 5 μg of Tmab where there was no treatment of cells. Lane 3 is cell lysate with no pull down. Lane 4 is the pull down using beads only. The ladder in kilodaltons (kDa) and is shown on the left.

### Bait-prey interaction models and identification of Accum-specific interactors

Due to the multiple components of ADCs and in particular Accum-T-DM1, we created two complimentary bait-prey interaction models to determine the Accum-specific interactors by Significance Analysis of INTeractome (SAINT). Tmab and T-DM1 were chosen as key references for the differential elimination of nonspecific prey proteins against the antibody backbone and drug DM1, respectively. This was named the 1 bait:3 control (I:III) model (Fig. 5A). Moreover, since Tmab is identical to the mAb in the experiment, it also served as the isotype control. Beads only controls were utilized, as they are a traditional negative reference for pull down experiments. Again a 2 h incubation period was justified as cytotoxicity effects were not yet observable such as rounding of the cells and detachment from the cell culture plate. Because this proteomic analysis of a NLS-modified agent for evaluating nuclear transport has never been performed according to our knowledge, we identified interactors based on a primary (high-confidence) and a secondary (medium-confidence) SAINT score (STSc) thresholds that were set at ≥0.9 and ≥0.5, respectively. The statistics are summarized in Suppl. Table S1. Using the cut-off threshold STSc≥0.9, only a single NTR, importin-β (KPNB1) was present. This was uncharacteristic since importin-β requires importin-α for NLS signal recognition and binding of cargoes destined for the nucleus. This suggested that the I:III model used in combination with an STSc ≥ 0.9 was overly stringent to identify potential NTR interactors. In contrast, using the cut-off of STSc ≥ 0.5, the specific interactors for Accum-T-DM1 included the NTRs importin 7 (IPO7), importin 4 (IPO4), importin-β, importin 5 (IPO5), transportin 1 (TNPO1), and importin-α (KPNA2).

**Figure 5.**
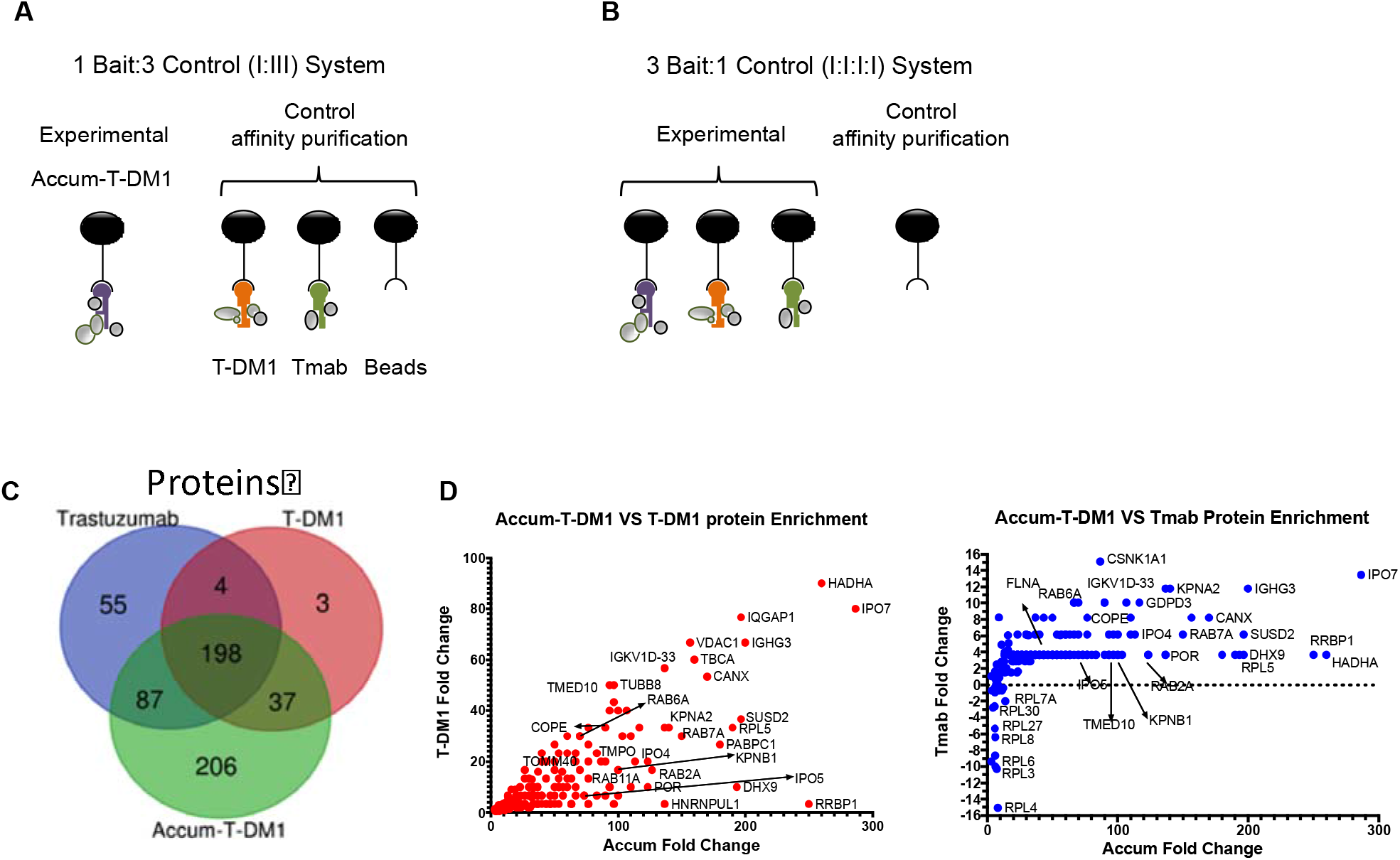
Interaction setup for Accum-T-DM1 as a (A) single bait with multiple controls (I:III) or (B) Accum-T-DM1, T-DM1, and Tmab as independent baits and a single common control (I:I:I:I). (C) I:I:I:I Venn diagram of enriched interactors. (D) I:I:I:I Prey enrichment comparison between Accum-T-DM1 versus T-DM1 (left) or Tmab (right).

For the second model, we performed SAINT analyses with Accum-T-DM1, T-DM1, and Tmab as individual baits and using beads only as the single control herein referred to as the I:I:I:I model (Fig. 3B). An added advantage of the second model was that interactors could be identified that were common and unique between the three baits. As the I:I:I:I model had reduced stringency based on controls, we considered only interactions with STSc≥0.9. As a result, there were 528, 242, and 347 proteins identified as interactors for Accum-T-DM1, T-DM1, and Tmab, respectively (Fig. 5C and Suppl. Table S1). Tmab and T-DM1 had 55 and three unique interactors, respectively. In contrast, Accum-T-DM1 had 206 unique interactors. There were 198 proteins common for all the baits. The large overlap was most likely due to the mAb Tmab, the only common feature shared by all three baits. Using the I:I:I:I model, there was an abundance of proteins with 10-300-fold enrichment for Accum-T-DM1 relative to T-DM1 and Tmab (Fig. 5D). Importantly, the exact number and type of NTR identified in model I:III at STSc≥0.5 were also significantly enriched for Accum-T-DM1 in model I:I:I:I at STSc≥0.9 (Fig. 5D). Thus, both models provided results in agreement with each other and together strengthen our certainty that the identified NTRs were specific Accum interactors.

With this novel prey-bait proteomic method it was also possible to characterize the expanding intracellular interaction network for Accum-T-DM1. Because Accum has a stretch of positively-charged amino acids we evaluated important characterizations as performed by Lang et al (*37*). As polylysine tagged proteins can present an experimental challenge for controlling nonspecific electrostatic interactions, molecular weight of interactors can be plotted and analyzed. We found that the majority of Accum-specific interactors had molecular weights of 10-30 kDa and there was a depletion of larger proteins (Suppl. Fig. S3A). Proteins between 10-30 kDa accounted for 50% of the Accum-specific interactors. A box plot analysis demonstrated there was a significant difference between distributions of molecular weight of Accum interactors and of all identified proteins (Suppl. Fig. S3B). Similar to the findings by Lang et al (*37*), enrichment of proteins with low molecular weights is typical of specific interactions and, hence, further supports that the identified interactome is Accum-specific.

### Molecular topology of protein interaction modules for Accum-T-DM1

The enriched interactors specific for Accum-T-DM1 from the SAINT analysis (I:III system) were entered into GeneMANIA (GM) in Cytoscape to determine enriched biological processes (Suppl. Table S3 and S4). Gene Set Enrichment Analysis (GSEA) was performed by GM to highlight the most enriched Gene Ontology (GO) terms and used to systematically attribute a ‘generic pathway’ to each protein in the network according to their main function (*38,39*). We identified 164 GO terms or pathways that were significantly enriched for Accum-T-DM1 (Suppl. Table S4). We identified 15 protein families or common cellular pathways that were significantly enriched in the dataset (Suppl. Table S3). Cytoscape allowed us to get an aerial overview of the interaction network by clustering proteins based on these protein families/pathways (Suppl. Fig. S4). The top five most enriched protein families/pathways were exclusively composed of ribosomes involved with unique cellular functions, including nuclear transport (Fig. 6 and Suppl. Table S4).

**Figure 6.**
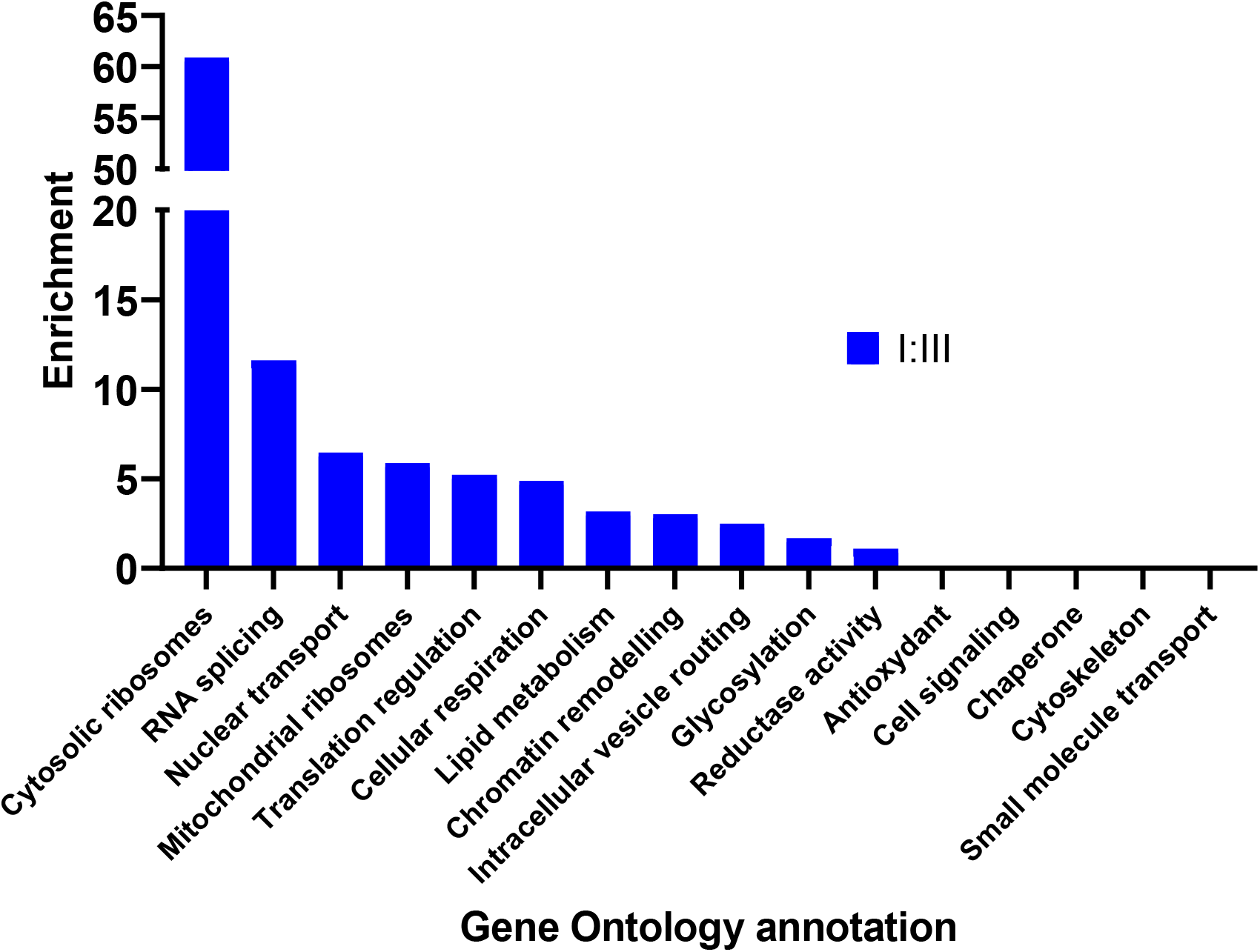
Enrichment values of a selection of GO annotation in the I:III system represented as a log_10_(1/q-value). Lipid metabolism, chromatin remodeling, intracellular vesicle routing and reductase activity are represented by energy derivation by oxidation of organic compounds, DNA geometric change, establishment of vesicle localization and NADH dehydrogenase complex respectively. Antioxidant, cell signaling, chaperone, cytoskeleton and small molecule transport did not match any representative GO terms in our analysis.

For the I:I:I:I model, genes obtained from the SAINT analysis with an STSc≥0.9 (Fig. 5B-D) were entered into GM (Suppl. Table S5 and S6). There were 310 GO terms identified for Accum-T-DM1, 231 for T-DM1 and 271 for Tmab. Of these, 192 were common between all three baits and 73 were unique to Accum-T-DM1 (Fig. 7A). The top five pathways represented on Figure 6 were also strongly enriched using the I:I:I:I model (Fig. 7B). Importantly, only Accum-T-DM1 had the *protein import into nucleus* GO term enriched. Thus, the I:III and I:I:I:I models generated overlapping data further confirming that this analytic proteomic method was specific for Accum.

**Figure 7.**
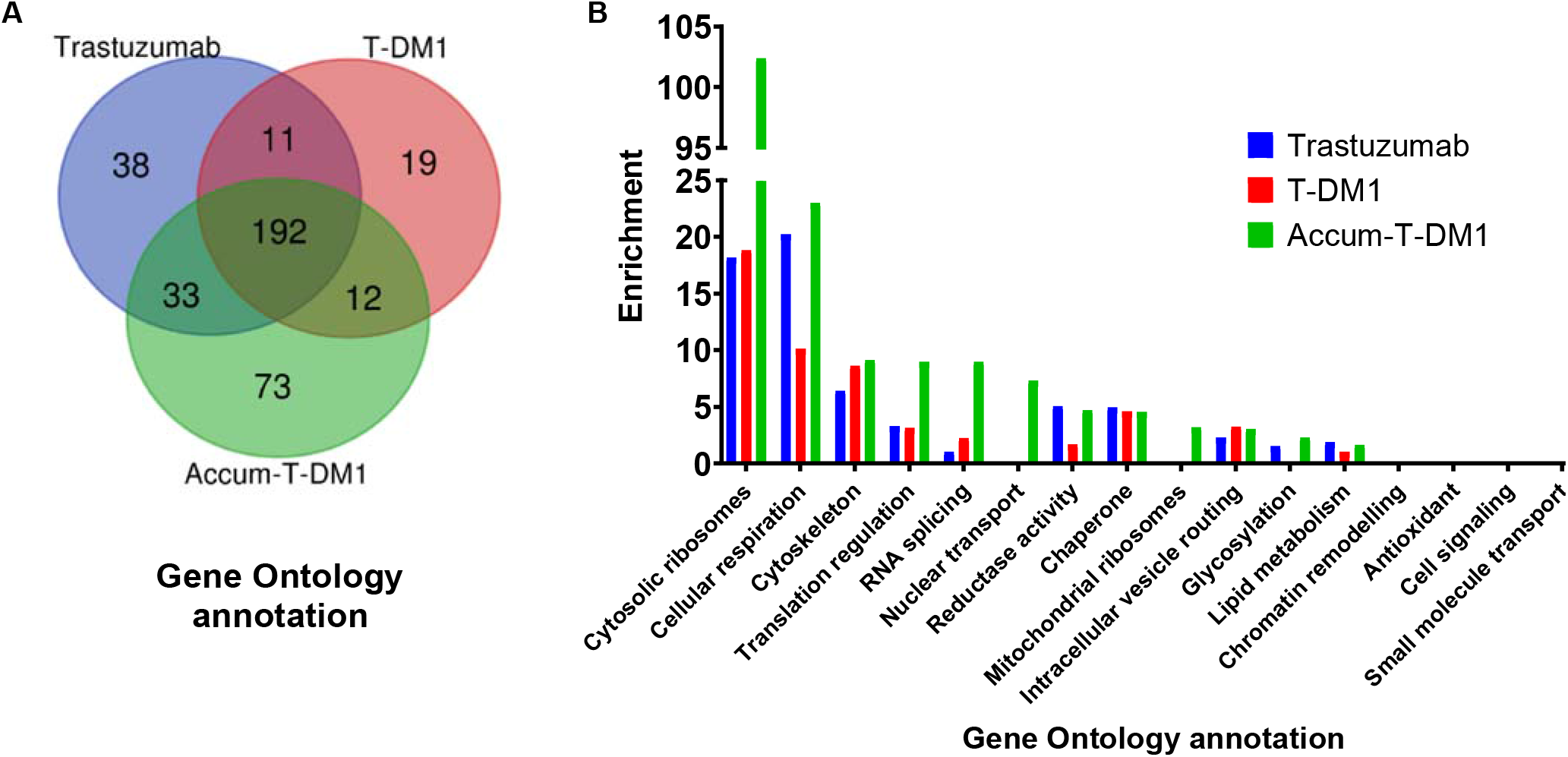
(A) Venn diagram of the enriched GO terms per bait in the I:I:I:I system. (B) Enrichment values of a selection of GO annotation in the I:I:I:I system represented as a log_10_(1/q-value). Cytoskeleton, reductase activity, chaperone, intracellular vesicle routing, and lipid metabolism are respectively represented by the GO terms microtubule, NADH dehydrogenase activity, response to unfolded protein, establishment of vesicle localization and lipid oxidation.

### Accum-T-DM1 NTR interactors

The affinity purification after cell treatment revealed unanticipated findings for NTRs interacting with Accum-T-DM1. As anticipated, the classical NTRs importin-α and importin-β had interaction values for Accum-T-DM1 above the STSc≤0.5 cut-off (Fig. 8A). However, the non-classical NTRs importin 4, importin 5, importin 7, and transportin 1 also had strong interaction values with Accum-T-DM1 (Fig. 8A). In fact, the non-classical NTR importin 7 was the most abundant off all NTRs with an average spectral count (AvgSpec) of 28.67 (STSc = 0.66). NTRs below the 0.5 Saint score threshold included, importin β3, importin 8 (IPO8), importin 9 (IPO9), importin 11 (IPO11), and karyopherin subunit α4 (KPNA4). The Saint scores, average spectral count, and Bayesian false discovery rate (BFDR) are listed in Supplemental Table S7.

**Figure 8.**
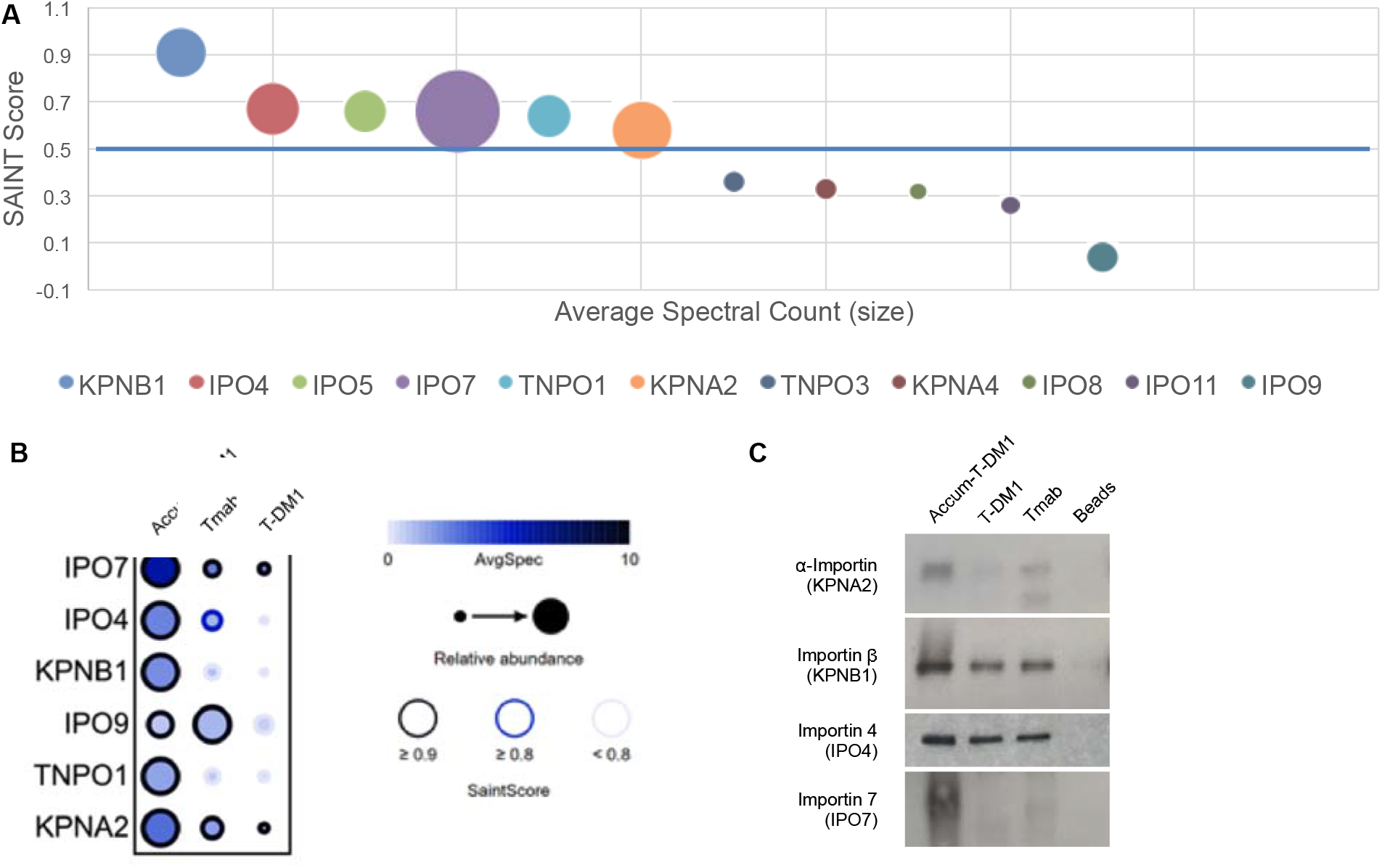
(A) Bubble plot showing the Saint score enrichment and overall abundance (dot size) of the identified NTRs. Cut off indicated by blue line. (B) Dot plot output showing quantitative SAINT interactions between Accum-T-DM1, Tmab, and T-DM1 and the NTRs found in the I:I:I:I system. The confidence of the detected interaction is shown as the circle edge, with black being high confidence (StSc ≥0.9), blue medium confidence (StSc ≥0.8), and light gray low confidence (StSc <0.8). Circle size indicates the prey’s relative abundance across baits. The bait that yielded the highest spectral count for a prey is given a full-sized circle, and the circle size for the other baits is scaled relative to this maximum. The color scale of the circles indicates the prey’s spectral count for the corresponding bait. (C) Confirmation of the NTR-Accum interaction by affinity purification followed by Western Blot against the indicated preys.

Figure 8B shows a dot plot generated from the I:I:I:I model to further demonstrate the specificity of enriched NTRs for Accum-T-DM1. The dot plot shows that importin 7 had a strong increase in relative abundance and average spectral count for Accum-T-DM1 relative to T-DM1, and Tmab. All NTRs with the exception of importin 9 with an STSc≥0.9 from Figure 8A were enriched for Accum-T-DM1 relative to Tmab and T-DM1. As the STSc was ≥0.9 for all three baits in the dot plot, the abundance of the NTR enriched is the defining factor and importin 7 is the most abundant.

Western blot was used to biochemically confirm the abundance of protein in the pull down assays of Accum-T-DM1, T-DM1, Tmab, and beads only in SKBR3 cells. In the Accum-T-DM1 pull down, importin 7 was the most abundantly present NTR (Fig. 8C). This validated the findings shown in the dot plot in Figure 8B. In addition, the presence of importin 7 was restricted to Accum-T-DM1 and was not present for T-DM1, Tmab, or the bead only pull down assays. For the classical NTRs, the presence of importin-β was increased relative to importin-α, which also confirmed the proteomic data. Importin-α and importin-β were also present in the T-DM1 and Tmab pull downs, albeit at reduced amounts compared to Accum-T-DM1. Importin 4 appeared to be only slightly less abundant in the T-DM1 and Tmab pull downs compared to Accum-T-DM1. This matched well with the dot plot as importin 4 had only a slightly increased average spectral count compared to Tmab. Thus, Western blot validated the findings from the proteomic data obtained from the I:I:I:I model.

### The requirement of the identified NTRs on Accum-T-DM1 cytotoxic effectiveness

The ability to use RNA interference as a tool for gene silencing was utilized to determine if an NTR identified in this proteomic method was essential to promote cytotoxic effectiveness for Accum-T-DM1. This would serve as a surrogate and key determinant to demonstrate the mechanistic and dependent link between the observed nuclear localization and an increased sensitivity of SKBR3 cells to Accum-T-DM1. SKBR3 cells were transfected with siRNA pools individually targeting KPNA2, KPNβ1, IPO4, IPO7, and siRNA control followed by lysing cells at 24 h and 96 h post transfection. Expression of indicated gene products targeted for silencing was evaluated at 24 h and 96 h posttransfection to coincide with the initiation and termination of the 72 h cytotoxicity assay of SKBR3 cells with T-DM1 and Accum-T-DM1. Western blots confirmed the reduction in protein levels for all four NTRs at 96 h (Fig.9A) but not at 24 h (Suppl. Figure S5) post transfection. Transfections with scrambled pools did not cause a reduction in NTR protein expression (Fig. 9A).

**Figure 9.**
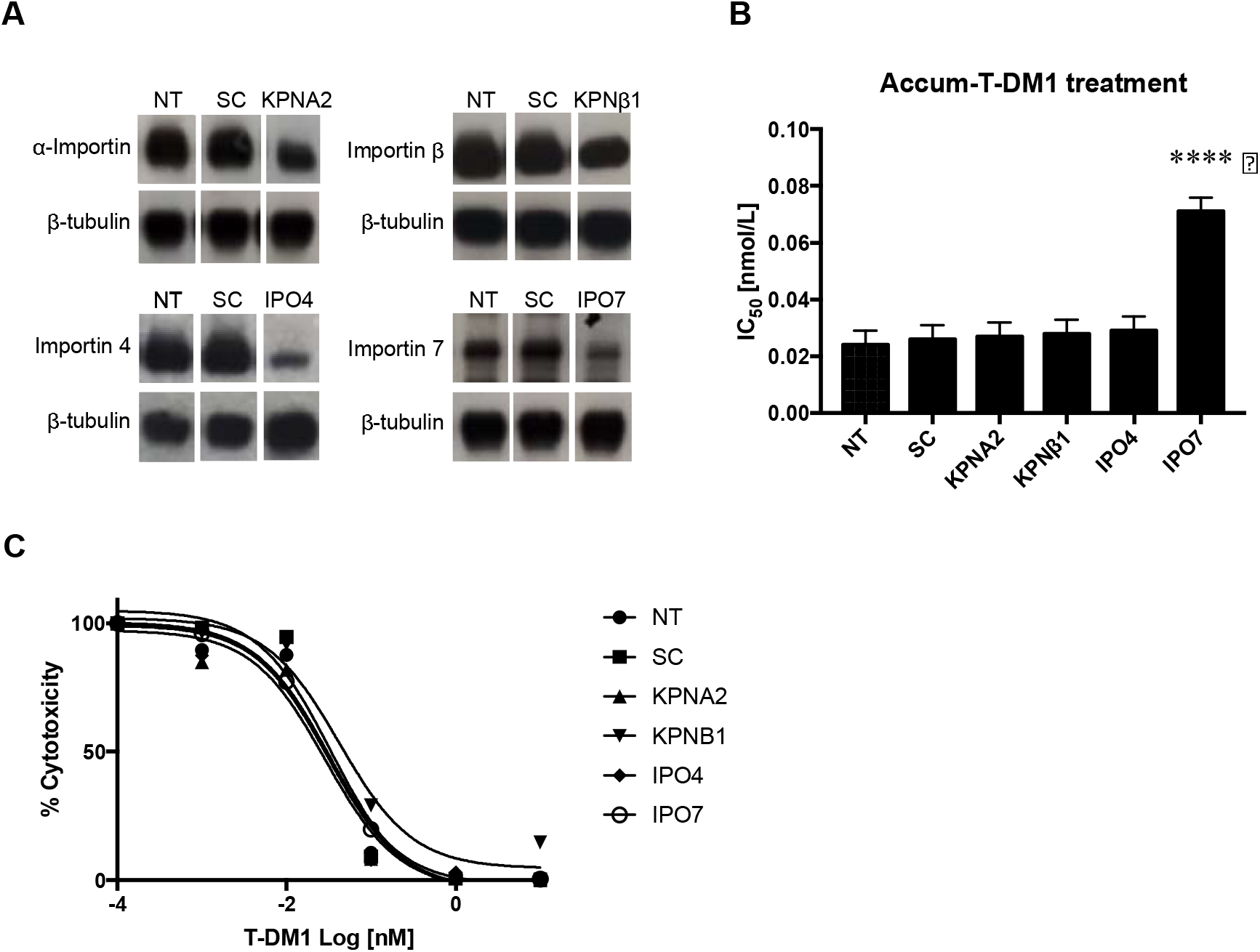
(A) Silencing of KPNA2, KPNβ1, IPO4, and IPO7 genes at 96 h post transfection and the Western blot images of the corresponding proteins α-importin, importin-β, importin 4, and importin 7. β-tubulin was used for the loading control. NT = not transfected, SC = Scrambled siRNA. (B) The IC_50_ values of Accum-T-DM1 after 72 h treatment of non-transfected (NT) SKBR3 cells and cells transfected with siRNA pools specific for indicated NTR genes or scrambled (SC) sequence. **** corresponds to p<0.0005. (C) SKBR3 cells treated 24 h after transfection with siRNA pools specific for indicated NTR genes, SC sequence or NT for 72 h with increasing concentrations of T-DM1

Accum-T-DM1 cytotoxicity was dependent on non-classical nuclear transport. The IC_50_ values for Accum-T-DM1 (with six Accum moieties [Accum_6_-T-DM1] on SKBR3 cells transfected with siRNA pools to knockdown KPNA2, KPNβ1, IPO4, and IPO7 gene expression were 0.027 nmol/L, 0.027 nmol/L, 0.029 nmol/L, and 0.071 nmol/L, respectively (Fig.9B). This result indicated that Accum-T-DM1 had a 3-fold loss in cytotoxic potency in SKBR3 cells when IPO7 was knocked down relative to the IC_50_ value of 0.024 nM with Accum_6_-T-DM1 in normal SKBR3 cells (Fig. 3A). In contrast, knockdown of the classical NTRs importin-α and importin-β did not affect the ability of Accum-T-DM1 to kill SKBR3 cells. The IC_50_ value of Accum-T-DM1 on SKBR3 cells transfected with scrambled siRNA pools was 0.026 nmol/L and further assured that the experiment was specific for IPO7. To assess specificity of IPO7-mediated transport of Accum-T-DM1, NTR-depleted SKBR3 cells were treated with increasing concentrations of standard T-DM1. There were no significant differences in IC_50_ values for all NTR knockdowns (Fig. 9C), indicating that IPO7 was the key NTR regulator for Accum-mediated shuttling of T-DM1 to the nucleus and cytotoxicity in SKBR3 cells.

## DISCUSSION

We developed an approach that was initially designed to confirm an established mechanism for nuclear transport of an agent harbouring the classical NLS from SV40 large T-antigen. Instead our results have opened interesting questions for researchers developing tools to study nuclear transport or for the development of NLS-based biopharmaceutical agents. Although, evidence of nuclear transport is showing that NTRs can be selective for endogenous cellular cargos depending on cell type or cellular states, NLS-modified agents continue to operate on a ‘hit and miss’ approach, particularly for therapeutic development. In addition, fundamental insights on nuclear transport using NLS-modified agents have been largely performed using cellular reconstituted systems. Thus, improved approaches that can dually improve rational therapeutic design of NLS-based agents and also acquire cellular insights due to nuclear transport in live cells are required.

In this study, we demonstrated that the Accum platform when conjugated to the clinically approved anti-HER2 biopharmaceutical T-DM1 was able to localize to the nucleus of SKBR3 cells and increase cytotoxicity by up to 18-fold (Fig. 3). We developed a generally applicable cellular treatment coupled to affinity pull down (Fig. 4) and proteomic method (Fig. 5A and B) for NLS-modified antibody-conjugate biopharmaceuticals. This allowed us to discover that the strongest NTR-specific interactor for Accum-T-DM1 was importin 7 (Fig. 8). The dependency of Accum-T-DM1 on importin 7 for its ability to improve tumor killing was demonstrated when its gene IPO7 was knocked down followed by treatment with Accum-T-DM1 and T-DM1 (Fig. 9A and B). The IC_50_ value in IPO7-knocked down cells treated with T-DM1 remained unchanged relative to non-knocked down cells (Fig. 9C). In contrast, the IC_50_ value for Accum-T-DM1 was decreased by 3-fold relative to the value obtained from the non-knocked down SKBR3 cytotoxicity assay with Accum-T-DM1 (Fig. 9B). Importantly, this suggested that the classical NTR complex of importin-α and importin-β was not required for nuclear transport and, hence, was the reason why knock down of the genes KPNA2 and KPNβ1 did not affect cytotoxicity of Accum-T-DM1.

Approximately one decade-ago Rix and Superti-Furga addressed the need by pharmaceutical communities to utilize proteomic profiling to better understand the interactions within cells of small molecules and has led to strong activity in this research domain (*40*). In contrast, studies on proteomic profiling of biopharmaceuticals with respect to their dynamics inside cells is scarce. Only recently has there been important advances that have utilized proteomic profiling to determine resistance mechanisms to biopharmaceuticals such as Tmab, T-DM1 and other emerging mAb-based drugs (*41–44*). However, these approaches have not specifically addressed intracellular transport. Mechanistic insights into intracellular transport of macromolecular therapeutics using proteomic approaches have only been reported for nanoengineered particles (*45,46*). These studies were essential as they elucidated key regulators in the intracellular transport of the nanoparticles. As a result, proteomic profiling of intracellular transport interactors has advanced the development of nanomaterials for improving the effectiveness of nanoengineered particles (*47*). Hence, this approach could also be adapted to provide improved effectiveness for NLS-modified agents as therapeutics.

It was important to develop a method whereby Accum-T-DM1, T-DM1, and Tmab were allowed to mix with cells at non-excessive concentrations and periods of time. The major challenge is to have sufficient amount of antibody for subsequent pulldowns. An added challenge was that pulldown efficiency was reduced due to Accum tagging to T-DM1 (Fig. 4). Thus, we expanded the number of cells per incubation to 15 x 10^6^ cells. In this work, our main goal was to not incubate cells with excessive amounts of drug or for periods that would lead to cytotoxicity, which could complicate our proteomic approach. As described, cells were checked to ensure cells were not undergoing apoptosis. Endo et al., used T-DM1 as bait to determine that DM1 binds a cell surface protein cytoskeleton-associated protein 5 (CKAP5) that is expressed on hepatocytes in the absence of HER2 (*48*). A concentration of 250 μg/mL (~1667 nmol/L) T-DM1 was incubated with 1.5 x 10^6^ cells. The study determined that DM1 interaction with CKAP5 is most likely a contributing factor for dose-limiting hepatotoxicity in patients treated with T-DM1. However, the T-DM1 was used at an extreme concentration. A direct comparison of cytotoxic potency between CKAP5- and HER2-positive only cells and T-DM1 was not reported. However, the potent cytotoxic concentration shown for T-DM1 in CKAP-positive/HER2-negative cells ranged from 30-to 142-fold higher than reported IC_50_ values for T-DM1 against HER2-positive cells (*27,33*), including this study. The method in our report is relatively benign, the cell number and drug concentration can can most likely be reduced, and can be generalized for additional NLS-tagged agents.

Although the term ‘nuclear transport’ can have different meanings between scientists studying basic mechanisms and those interested in therapy, the development of SV40 NLS-modified agents is an area of research that is actively investigated (*49–54*). However, the general consensus is the mechanism for these NLS-containing drugs is dependent on the importin-α/importin-β complex to achieve nuclear localization (*52–55*). Our results clearly indicate that the SV40 NLS as part of Accum is dependant on non-classical nuclear transport. Yet, the development of SV40 NLS-tagged agents for the development of various therapeutic agents is ongoing (*55–60*). In addition, different NLS sequence-tagged therapeutics have made significant advances. For example, HIV-1 trans-activating protein harbours an NLS and has been extensively utilized for the nuclear delivery of various agents (*61–63*), with a few reaching the clinic (NCT01975116, NCT00914914). However, they have been shown to suffer from target antigen specificity problems (*64,65*).

The additional interactors uncovered by the proteomic portion of this study provide further impact to future directions on studying nuclear transport using NLS-modified agents. A majority of the human non-classical NTRs are able to recognize unique sets of proteins or RNA, thus creating multiple transport pathways across the NPC (*18*). Relevant to this study, importin 7 is known to mediate nuclear transport by forming a heterodimer with importin-β. This supports why the strong interaction of importin-β with Accum-T-DM1 was continuously present in the different proteomic analyses used in this study. Because we showed that cytotoxicity was impaired only with importin 7, this suggests that if a complex exists in SKBR3 cells with importin-β and importin-7, transport of Accum-T-DM1 is only dependent on the presence of importin 7. Importin 7 is also known to be very important for the nuclear transport of ribosomal proteins and histones (*18*). Interestingly, histone 1 binds several NTRs but import was only achieved in the presence of both importin 7 and importin-β (*66,67*). However, in this study we showed the opposite. Accum-T-DM1 is reliant solely on importin 7 and not on importin-β. This suggests that in SKBR3 cells, in the context of an SV40 large T-antigen NLS-modified agent, transport is likely only mediated by importin 7 and not an importin 7/importin-β heterodimer. Preference for importin 7 versus an importin 7/importinβ heterodimer has been observed for particular nuclear localized proteins such as EZI and ERK-2 kinase (*68,69*). Importin 7 has also been shown to transport RBL 4 and RBL 6 into the nucleus (*67*), which were both identified to interact with Accum-T-DM1. Importin 4 that contained the second highest SAINT score among NTRs was not essential for Accum-T-DM1 cytotoxicity. Importin 4 is known to bind to only a few known proteins (*18*). Among them the protein TP2 contains an NLS sequence of GKVSKRKAV. Since this sequence is similar to the NLS sequence contained in Accum it is logical that importin 4 interacted with Accum-T-DM1. However, it was not essential for nuclear transport and cytotoxicity. Hence, only importin 7 can be expected to have potential impact on Accum’s ability to improve cytotoxicity for T-DM1 in SKBR3 cells. Moreover, Accum resembles ribosome and histone protein topology and, hence, appears to naturally incorporate into the nuclear transport mechanism involving importin 7.

This proteomic and biochemical approach can be adopted for any NLS-modified mAb agent – a foresight for rationale approach for studying nuclear transport mechanics or for the development of therapeutic agents (Fig. 10). Thus, this work sets the stage for future studies to determine a pattern for NTR reliance for NLS-modified agents among different cellular systems and contexts. For next-generation NLS-modified biopharmaceuticals, it is profoundly important to understand the tumor system to optimize the NLS-based approach. Such understandings could be useful in combination with DNA/RNA gene therapy approaches where the NTRs specific for ribosome nuclear transport could synergize to increase nuclear delivery efficiency. For fundamental insight, our approach can be combined with gene expression studies of various NTRs to understand the cellular response to treatment with NLS-modified agents. Thus, aspects such as agent concentrations and incubation time points can be further refined in order to delineate the most accurate transport network for a given agent. This method can be further exploited to select specific NLS sequences for particular NTRs that have preferential effects for a given application. Lastly, an advantage of our current methodology is that these measurements can be made in a structured manner using easy *in vitro* biochemical methods and is combined with a proteomic system that can be easily applied to any NLS-modified agent.

**Figure 10.**
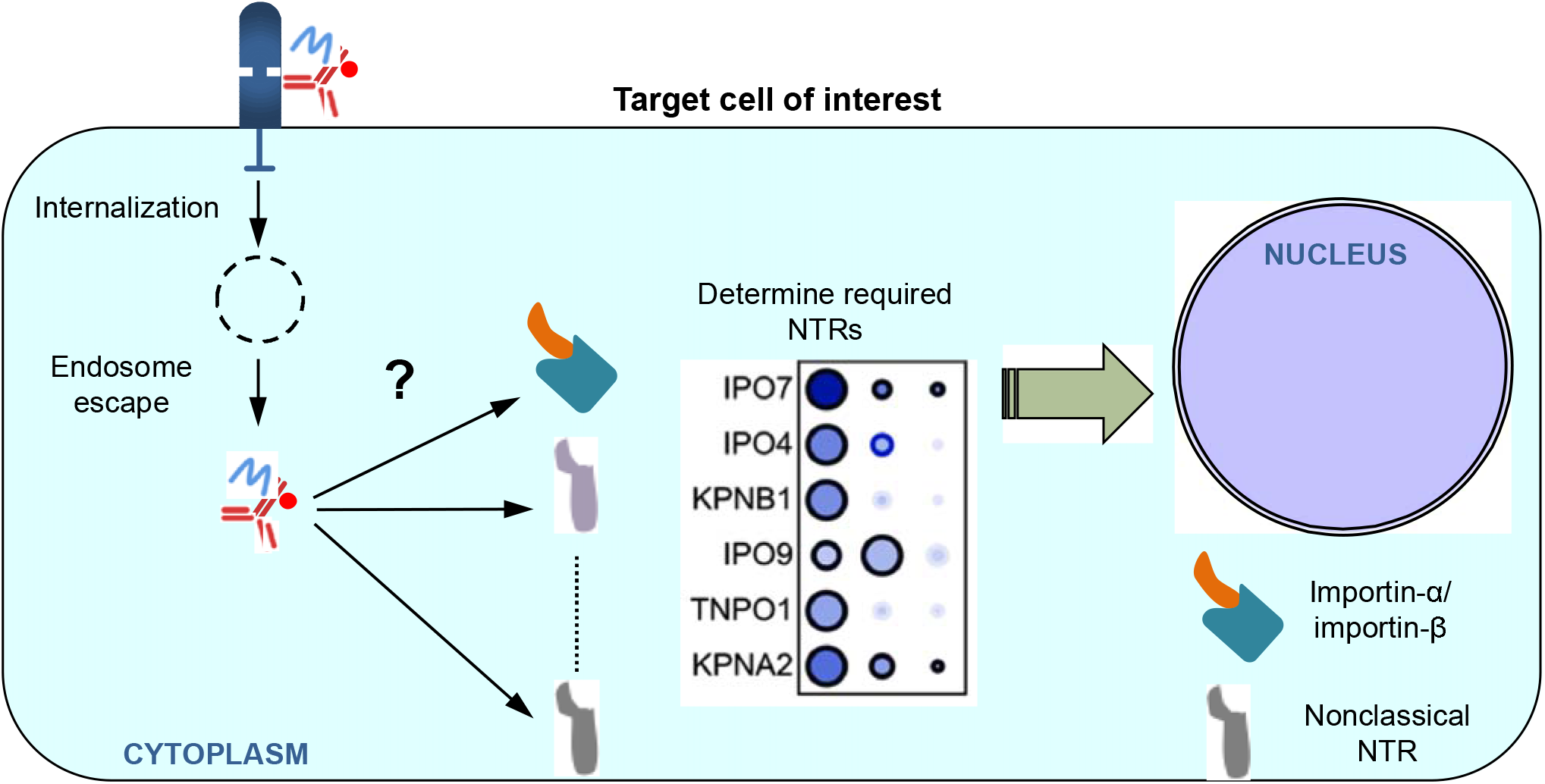
A foresight view of an NLS-modified mAb agent that binds its specific receptor on the cell surface and is internalized. In the study, Accum is able to escape endosome entrapment, The approach is to then lyse cells and pull down the mAb and conduct a proteomic analysis to determine the required NTRs for nuclear transportation and effectiveness of the agent.

## MATERIALS AND METHODS

### Cell culture

SKBR3 and MCF7 cells were obtained from ATCC and were tested for authenticity and contamination with viruses or mycoplasma prior to experimentation. Cells were grown in accordance with ATCC recommendations.

### Accum conjugation

Accum was synthesized as previously described (*22*). T-DM1 was obtained from the CHUS pharmacy. The SM(PEG)_2_ was reacted in molar excess to 200 ⌈g of T-DM1 in order to obtain approximately different amounts of Accum moieties per T-DM1. Reaction conditions to control the amount of Accum per mAb have been previously described (*70*). Accum-modified T-DM1 was then transferred to a Centricon YM-100 ultrafiltration tube (EMD Millipore, Ontario, Canada) and concentrated in PBS, pH 7.4. 10 ⌈g of T-DM1 and Accum-T-DM1 ADCs were loaded on to a 12% polyacrylamide gel. For determination and calculation of Accum moieties per T-DM1 molecule, conjugates were analyzed by SDS-PAGE under reducing conditions on a 12% Tris-HCl polyacrylamide gel and stained with Coomassie Brilliant Blue R-250 (BioRad, Ontario, Canada). The migration distance in the gel relative to the blue dye front (Rf) was measured and the number of Accum moieties introduced into the light chain and heavy chain of T-DM1 were estimated by reference to a logarithm plot of molecular weight versus 1/Rf for Kaleidoscope Prestained Standards (BioRad) electrophoresed under identical conditions.

### Differential Scanning Fluorimetry

Lyophilized T-DM1 was suspended in PBS and the Accum-T-DM1 formulations were evaluated from solution obtained after concentration. 10 ⌈L of 1 mg/mL ADCs were loaded into standard capillaries and mounted in a Prometheus NT.48 (Nanotemper Technologies, Germany) with excitation near UV. The temperature gradient was set to 1 °C/min in the range of 20 – 95 °C. ADC unfolding was measured by detecting the change in tryptophan/tyrosine fluorescence at emission wavelengths of 330 and 350 nm as a function of temperature. Melting temperatures were determined by detecting the maximum of the first derivative of the fluorescence ratios (350nm/330nm). Raw data was analyzed using the ThermControl software and statistics were calculated using Excel.

### Flow cytometry

1 x 10^6^ SKBR3 cells were seeded in 6-well plates 24 h prior to experimentation. Cells were washed once with PBS and then treated with 7.5μg/mL of T-DM1 or Accum-T-DM1 in media for 15 minutes (min), 30 min, 1 hour (h), 2 h, 4 h, 6 h, and 8 h. At the end of each indicated time, cells were lifted with 250 μl of 0.25% trypsin/ETDA (Wisent) for 5 min at RT, suspended in 1 ml of complete media and centrifuged for 5 min a 1000 x g. Supernatant was removed and cells were washed twice with ice cold PBS. Washed cells were then fixed in 1% paraformaldehyde (PFA) and 1% sucrose in PBS for 30 min on ice. Fixed cells were then washed twice with PBS and permeabilized using 0.015% Triton X-100 in PBS for 10 min at room temperature (RT). Cells were then incubated a rabbit anti-human IgG conjugated to Alexa Fluor 647 (Invitrogen) for 1 h, on ice, in the dark, washed twice in PBS. 10 μg/ml of propidium iodine (PI) was then added to the tubes. Fluorescence was measured on a Cytoflex 15 (Beckman Coulter) and 10,000 cells per sample were analyzed. Data was analyzed using CytExpert (version 2.0, Beckman Coulter).

### Immunofluorescence confocal microscopy

SKBR3 cells in media were treated with 7.5 ug/mL of Accum-T-DM1 and T-DM1 for up to 2 h at 37 °C. Cells were fixed using 4% PFA and 4% sucrose in PBS at 4 °C for 30 min, and permeabilized with 0.1% Triton X-100 in PBS for 15 min at RT. Subsequent steps were made in the blocking buffer containing 2% bovine serum albumin, 0.1% Triton X-100 in PBS. Blocking was performed for 30 min at RT and incubation with rabbit anti-human AlexaFluor 647 (Invitrogen) that binds to the Tmab Fc region of T-DM1 and Accum-T-DM1. Nuclear staining was performed with Hoechst (1:10,000) concurrently for 1 h at RT. Cells were washed, processed, and images acquired as previously described (*22*).

### Cytotoxicity assays

5 x 10^3^ SKBR3 (normal and knocked down) cells in 96-well flat-bottom plates were exposed to increasing concentrations of T-DM1 and Accum-T-DM1 at 37 °C for 72 h. Media was removed and cells were incubated with PrestoBlue (ThermoFisher) for 20 min and measured in a fluorometric plate reader in accordance with the manufacturer protocol. Assays were performed in triplicate.

### SKBR3 cell treatment and affinity purification

15 x 10^6^ SKBR3 cells were seeded in a 150 mm dish 24 h prior to experimentation. Cells were treated with 7.5 μg/mL of Tmab, T-DM1 or Accum-T-DM1 for 2 h at 37 °C. Cells were then washed twice with PBS at RT and then lysed with ice-cold RIPA buffer (Tris-HCl 50 mM, pH 7.5 at 25 °C, NaCl 150 mM, 1% v/v Triton X-100, 0.5% w/v sodium deoxycholate, PMSF 1 mM, NaF 5 mM). Lysates were passed 10 times through a 21-gauge needle and tubes centrifuged at 20,000g for 5 min to pellet insoluble cell debris. Supernatants were transferred to fresh tubes and diluted at a 1:1 ratio in RIPA buffer. 25 μl of protein G-coated magnetic beads (Thermo Fisher Scientific, 10003D) were equilibrated by washing twice in RIPA buffer and then mixed with 1.5 mL of diluted cell lysate for 1 h at RT with inversion. Beads were isolated on a magnetic rack and washed four times in PBS. The pulled down proteins were then processed for high performance liquid chromatography (HPLC)-MS/MS analysis or Western blot.

### Sample Preparation for HPLC-MS/MS

Beads from pulldowns were transferred to fresh tubes and washed five times with 20 mM NH4HCO3 in MS grade water. After the final wash, the beads were suspended in 100 μl of NH4HCO3 buffer containing 10 mM dithiothreitol (DTT) and incubated at 60 °C with mixing for 30 min. Tubes were cooled to RT and 15 mM of iodoacetamide was added and incubated for an additional 1 h at RT in the dark with mixing. The iodoacetamide was then quenched by the addition of 1 M DTT to a final concentration of 15 mM DTT and left for 10 min.

On bead digestion was performed by adding 1 μg of MS-grade trypsin (Promega, V5280) to each tube and incubating at 37 °C with shaking overnight. The following day, MS-grade formic acid (FA) up to 1% final concentration was added to quench digestion. Tubes were set on a magnetic rack to pellet the beads and the supernatant containing the peptides were transferred to fresh tubes. The beads were then washed in 100 μl of 60% MS-grade acetonitrile in 0.1% FA for 5 min. The supernatants were then pooled and then dried under vacuum. Peptide desalting and HPLC-MS/MS was performed using the identical methods and parameters as described by Chauvin et al (*71*). The sample preparation and HPLC-MS/MS were performed five independent times using freshly obtained T-DM1 and prepared Accum-T-DM1. The mass spectrometry proteomics data have been deposited to the ProteomeXchange Consortium via the PRIDE partner repository (*72*) with the dataset identifier PXD014786.

### Label-free quantification data analysis

MS raw files from the performed HPLC-MS/MS were analyzed with MaxQuant software (version 1.6.0.1). Specificity was set to trypsin, defined as cleavage after a lysine or an arginine not before a proline, maximum of 2 missed cleavages allowed, and peptides had to be at least 7 amino acids long. Variable amino acid modification included methionine oxidation and protein N-terminal acetylation. Fixed modification included cysteine carbamidomethylation. The mass tolerance for precursor and fragment ions was 7 ppm, and 20 ppm, respectively. Spectra were searched against the UniprotKB (Homo sapiens, 11/12/2018, 88,354 entries (*73*)) for protein identification with a false discovery rate (FDR) of 1%. Potential contaminants, reverse peptides, proteins only identified by sites, with less than two unique peptides, and duplicated proteins in the data matrix were excluded. The three replicates that exhibited the most internal reproducibility, to avoid artificial enrichment or impoverishment due to variation in sample processing, were selected for further analysis. The resulting protein groups can be found in Supplemental Table S1.

### SAINT identification of Accum-specific interactors

SAINT determined the Accum-specific interactors. Spectral counts obtained from MaxQuant were imputed into CRAPome. SAINT analysis was performed on the I:III mode using STSc thresholds value set at ≥0.9 and ≥0.5, respectively. The I:I:I:I model was also analyzed by SAINT with a STSc cut-off of 0.9 and included CRAPome controls (CC405, 406, and 410) (*74*). SAINT parameters are available in Supplementary Tables S3 and S5 for the I:III and I:I:I:I models, respectively.

### Network Visualization and Gene Set Enrichment Analysis

For the I:III model, a gene list was extracted from the SAINT results using the STSc≥0.5 and entered into GM (version 3.5.0 (*75*)) in Cytoscape (version 3.6.1). The SAINT analysis results were then incorporated into Cytoscape. GSEAwas performed by GM to determine the most enriched GO terms. These GO terms were then used to systematically attribute a ‘generic pathway’ to each protein in the network according to their main function. Proteins that could not be attributed a generic pathway through GM were manually searched in Uniprot Knowledgebase (*73*), Human Protein Atlas (*76–78*) and Pubmed and assigned to their most significant function. The GM parameters, results and generic pathway annotations can be found in Supplemental Table S3. GO terms and q-values are listed in Supplemental Table S4.

For the I:I:I:I model, genes obtained from the SAINT analysis with an STSc≥0.9 were inputted into GM. The GSEA results were extracted and common pathways were assessed using the Bioinformatics & Evolutionary Genomics tool to generate a Venn diagram (http://bioinformatics.psb.ugent.be/webtools/Venn/). Q-values obtained from GM were transformed for visualization purpose with the following equation: log(1/q-value). The I:I:I:I model was used to validate the findings from the I:III model thus, generic pathways were only assigned to individual genes using GM analysis results (Suppl. Table S5), without manual search through databases. The list of GO terms and enrichment values for the baits are listed in Supplemental Table S6.

### Gene silencing

SiRNA (pool of 4) directed against the indicated genes were obtained from Dharmacon. 3.5 x 10^5^ cells were transfected according to the manufacturer protocol using DharmaFECT for 24 h in serum-free medium (Life Technologies) in 6-well plates. The cells were then split for cytotoxicity assays and Western blot analysis. Transfections were performed in triplicate for each condition. Cytotoxicity assays for gene-silenced cells was performed as previously described.

### Western blot

Western blot was performed to evaluate the MS-identified NTR interactors to determine gene knockdown. After washing in PBS, the beads were suspended in 50 μl of 2x loading buffer (100 mM Tris-HCl pH 6.8, 2% SDS, 12% glycerol, 10 mM DTT and 0.02% bromophenol blue) and boiled for five min. Beads were captured on a magnetic support and 25 μl of supernatant was loaded onto an 8% SDS gel and electrophoresed for 2 h at 150V. The bicinchoninic acid assay (Bio-Rad) was used to measure protein concentration from the cell lysates and 20 μg of protein was loaded onto 8% SDS gels and electrophoresed. Proteins were then transferred to a polyvinylidene difluoride (PVDF) membrane overnight, at 4°C under a constant current of 40 mA on ice. PVDF membranes were then rinsed three times in H_2_O, dried to fix proteins in place, rehydrated in 100% methanol, washed twice in PBS, twice in Tris-HCl 50mM pH 7.5, NaCl 150mM, tween 20 0,1% v/v (TBST) and blocked for 1 h in 5% fat-free milk in TBST with gentle shaking. Membranes were then washed three times with TBST and incubated with mouse polyclonal anti-IPO7 (1:200), rabbit polyclonal anti-IPO4 (1:400), rabbit anti-KPNB1 (1:400), or goat polyclonal anti-KPNA2 (1:500). The mouse polyclonal anti-tubulin (1:500) was used for loading controls. Anti-IPO7, IPO4, and KPNB1 antibodies were purchased from LifeSpan Bioscience. Anti-KPNA2 and tubulin antibodies were purchased from Invitrogen. Incubations were performed at 4 °C, in 2.5% milk in TBST, with gentle shaking overnight. Membranes were then washed in TBST three times and incubated for 1 h, at RT in 2.5% milk in TBST with HRP-conjugated goat anti-rabbit (1:4000), goat anti-mouse (1:5000) (Invitrogen), or donkey anti-goat (1:5000) (Cedarlane). Finally, blots were washed twice with TBST, twice with PBS and revealed by electrochemiluminescence (Thermo Fisher Scientific).

### Statistics

Significance testing for comparisons between different IC_50_ values was performed using a 1-way ANOVA with Turkey’s multiple comparisons test.

## ACKNOWLEDGEMENTS

This work was supported by funds from the Canadian Institutes of Health Research and Fonds de recherche Quebec – Santé (FRQS). Simon Barsalou for conducting immunofluorescent confocal experiments. Thank you to the proteomic platform at the FMSS. S.J. and J.V.L are members of the FRQS-funded Centre de recherche du CHUS. S.J. and J.V.L also hold faculty salary awards from FRQS. Dr. Myriam Badr for discussions about stability data.

## Author Contributions

Conception and design: V. L. prepared and characterized ADCs, designed the proteomic approach, performed all experiments, and co-wrote manuscript. S.B and S.J helped conceptualize experiments and to interpret data. J.V.L conceptualized and directed the overall approach and co-wrote the manuscript.

## REFERENCES

1. Hobernik D, Bros M. DNA Vaccines-How Far From Clinical Use? Int J Mol Sci. 2018;19.

2. Bavelaar BM, Lee BQ, Gill MR, Falzone N, Vallis KA. Subcellular Targeting of Theranostic Radionuclides. Front Pharmacol. 2018;9:996.

3. Edens LJ, White KH, Jevtic P, Li X, Levy DL. Nuclear size regulation: from single cells to development and disease. Trends Cell Biol. 2013;23:151–159.

4. Loveland KL, Major AT, Butler R, Young JC, Jans DA, Miyamoto Y. Putting things in place for fertilization: discovering roles for importin proteins in cell fate and spermatogenesis. Asian J Androl. 2015;17:537–544.

5. Fried H, Kutay U. Nucleocytoplasmic transport: taking an inventory. Cell Mol Life Sci. 2003;60:1659–1688.

6. Macara IG. Transport into and out of the nucleus. Microbiol Mol Biol Rev. 2001;65:570–594, table of contents.

7. Brownlee C, Heald R. Importin alpha Partitioning to the Plasma Membrane Regulates Intracellular Scaling. Cell. 2019;176:805–815 e808.

8. Levy DL, Heald R. Nuclear size is regulated by importin alpha and Ntf2 in Xenopus. Cell. 2010;143:288–298.

9. Yasuda Y, Miyamoto Y, Yamashiro T, et al. Nuclear retention of importin alpha coordinates cell fate through changes in gene expression. EMBO J. 2012;31:83–94.

10. Yamaguchi YL, Tanaka SS, Yasuda K, Matsui Y, Tam PP. Stage-specific Importin13 activity influences meiosis of germ cells in the mouse. Dev Biol. 2006;297:350–360.

11. Li KK, Yang L, Pang JC, et al. MIR-137 suppresses growth and invasion, is downregulated in oligodendroglial tumors and targets CSE1L. Brain Pathol. 2013;23:426–439.

12. Szczyrba J, Nolte E, Hart M, et al. Identification of ZNF217, hnRNP-K, VEGF-A and IPO7 as targets for microRNAs that are downregulated in prostate carcinoma. Int J Cancer. 2013;132:775–784.

13. Wang CI, Chien KY, Wang CL, et al. Quantitative proteomics reveals regulation of karyopherin subunit alpha-2 (KPNA2) and its potential novel cargo proteins in nonsmall cell lung cancer. Mol Cell Proteomics. 2012;11: 1105–1122.

14. Adam SA, Gerace L. Cytosolic proteins that specifically bind nuclear location signals are receptors for nuclear import. Cell. 1991;66:837–847.

15. Gorlich D, Kutay U. Transport between the cell nucleus and the cytoplasm. Annu Rev Cell Dev Biol. 1999;15:607–660.

16. Kalderon D, Roberts BL, Richardson WD, Smith AE. A short amino acid sequence able to specify nuclear location. Cell. 1984;39:499–509.

17. Lanford RE, Butel JS. Construction and characterization of an SV40 mutant defective in nuclear transport of T antigen. Cell. 1984;37:801–813.

18. Chook YM, Suel KE. Nuclear import by karyopherin-betas: recognition and inhibition. Biochim Biophys Acta. 2011;1813:1593–1606.

19. Jakel S, Gorlich D. Importin beta, transportin, RanBP5 and RanBP7 mediate nuclear import of ribosomal proteins in mammalian cells. EMBO J. 1998;17:4491–4502.

20. Mosammaparast N, Jackson KR, Guo Y, et al. Nuclear import of histone H2A and H2B is mediated by a network of karyopherins. J Cell Biol. 2001;153:251–262.

21. Muhlhausser P, Muller EC, Otto A, Kutay U. Multiple pathways contribute to nuclear import of core histones. EMBO Rep. 2001;2:690–696.

22. Beaudoin S, Rondeau A, Martel O, Bonin MA, van Lier JE, Leyton JV. ChAcNLS, a Novel Modification to Antibody-Conjugates Permitting Target Cell-Specific Endosomal Escape, Localization to the Nucleus, and Enhanced Total Intracellular Accumulation. Mol Pharm. 2016;13:1915–1926.

23. Paquette M, Vilera-Perez LG, Beaudoin S, et al. Targeting IL-5Ralpha with antibody-conjugates reveals a strategy for imaging and therapy for invasive bladder cancer. Oncoimmunology. 2017;6:e1331195.

24. Paquette M, Beaudoin S, Tremblay MA, et al. NLS-Cholic Acid Conjugation to IL-5Ralpha-Specific Antibody Improves Cellular Accumulation and In Vivo Tumor-Targeting Properties in a Bladder Cancer Model. Bioconjug Chem. 2018;29:1352–1363.

25. Ritchie M, Tchistiakova L, Scott N. Implications of receptor-mediated endocytosis and intracellular trafficking dynamics in the development of antibody drug conjugates. MAbs. 2013;5:13–21.

26. Granger E, McNee G, Allan V, Woodman P. The role of the cytoskeleton and molecular motors in endosomal dynamics. Semin Cell Dev Biol. 2014;31:20–29.

27. Lewis Phillips GD, Li G, Dugger DL, et al. Targeting HER2-positive breast cancer with trastuzumab-DM1, an antibody-cytotoxic drug conjugate. Cancer Res. 2008;68:9280–9290.

28. Oroudjev E, Lopus M, Wilson L, et al. Maytansinoid-antibody conjugates induce mitotic arrest by suppressing microtubule dynamic instability. Mol Cancer Ther. 2010;9:2700–2713.

29. Wakankar AA, Feeney MB, Rivera J, et al. Physicochemical stability of the antibody-drug conjugate Trastuzumab-DM1: changes due to modification and conjugation processes. Bioconjug Chem. 2010;21:1588–1595.

30. Wang L, Amphlett G, Blattler WA, Lambert JM, Zhang W. Structural characterization of the maytansinoid-monoclonal antibody immunoconjugate, huN901-DM1, by mass spectrometry. Protein Sci. 2005;14:2436–2446.

31. Buecheler JW, Winzer M, Tonillo J, Weber C, Gieseler H. Impact of Payload Hydrophobicity on the Stability of Antibody-Drug Conjugates. Mol Pharm. 2018;15:2656–2664.

32. Luo Q, Chung HH, Borths C, et al. Structural Characterization of a Monoclonal Antibody-Maytansinoid Immunoconjugate. Anal Chem. 2016;88:695–702.

33. Junttila TT, Li G, Parsons K, Phillips GL, Sliwkowski MX. Trastuzumab-DM1 (T-DM1) retains all the mechanisms of action of trastuzumab and efficiently inhibits growth of lapatinib insensitive breast cancer. Breast Cancer Res Treat. 2011;128:347–356.

34. Subik K, Lee JF, Baxter L, et al. The Expression Patterns of ER, PR, HER2, CK5/6, EGFR, Ki-67 and AR by Immunohistochemical Analysis in Breast Cancer Cell Lines. Breast Cancer (Auckl). 2010;4:35–41.

35. Austin CD, De Maziere AM, Pisacane PI, et al. Endocytosis and sorting of ErbB2 and the site of action of cancer therapeutics trastuzumab and geldanamycin. Mol Biol Cell. 2004;15:5268–5282.

36. Ram S, Kim D, Ober RJ, Ward ES. The level of HER2 expression is a predictor of antibody-HER2 trafficking behavior in cancer cells. MAbs. 2014;6:1211–1219.

37. Lang WH, Calloni G, Vabulas RM. Polylysine is a Proteostasis Network-Engaging Structural Determinant. J Proteome Res. 2018;17:1967–1977.

38. Ashburner M, Ball CA, Blake JA, et al. Gene ontology: tool for the unification of biology. The Gene Ontology Consortium. Nat Genet. 2000;25:25–29.

39. The Gene Ontology C. The Gene Ontology Resource: 20 years and still GOing strong. Nucleic Acids Res. 2019;47:D330–D338.

40. Rix U, Superti-Furga G. Target profiling of small molecules by chemical proteomics. Nat Chem Biol. 2009;5:616–624.

41. Liu W, Chang J, Liu M, et al. Quantitative proteomics profiling reveals activation of mTOR pathway in trastuzumab resistance. Oncotarget. 2017;8:45793–45806.

42. Loganzo F, Tan X, Sung M, et al. Tumor cells chronically treated with a trastuzumab-maytansinoid antibody-drug conjugate develop varied resistance mechanisms but respond to alternate treatments. Mol Cancer Ther. 2015;14:952–963.

43. Ramello MC, Benzaid I, Kuenzi BM, et al. An immunoproteomic approach to characterize the CAR interactome and signalosome. Sci Signal. 2019;12.

44. Sung M, Tan X, Lu B, et al. Caveolae-Mediated Endocytosis as a Novel Mechanism of Resistance to Trastuzumab Emtansine (T-DM1). Mol Cancer Ther. 2018;17:243–253.

45. Hofmann D, Tenzer S, Bannwarth MB, et al. Mass spectrometry and imaging analysis of nanoparticle-containing vesicles provide a mechanistic insight into cellular trafficking. ACS Nano. 2014;8:10077–10088.

46. Yan Y, Lai ZW, Goode RJ, et al. Particles on the move: intracellular trafficking and asymmetric mitotic partitioning of nanoporous polymer particles. ACS Nano. 2013;7:5558–5567.

47. Park JS, Yi SW, Kim HJ, et al. Verification of Long-Term Genetic Stability of hMSCs during Subculture after Internalization of Sunflower-Type Nanoparticles (SF-NPs). Theranostics. 2018;8:5548–5561.

48. Endo Y, Takeda K, Mohan N, et al. Payload of T-DM1 binds to cell surface cytoskeleton-associated protein 5 to mediate cytotoxicity of hepatocytes. Oncotarget. 2018;9:37200–37215.

49. Christie M, Chang CW, Rona G, et al. Structural Biology and Regulation of Protein Import into the Nucleus. J Mol Biol. 2016;428:2060–2090.

50. Escriou V, Carriere M, Scherman D, Wils P. NLS bioconjugates for targeting therapeutic genes to the nucleus. Adv Drug Deliv Rev. 2003;55:295–306.

51. Pichon C, Billiet L, Midoux P. Chemical vectors for gene delivery: uptake and intracellular trafficking. Curr Opin Biotechnol. 2010;21:640–645.

52. Pouton CW, Wagstaff KM, Roth DM, Moseley GW, Jans DA. Targeted delivery to the nucleus. Adv Drug Deliv Rev. 2007;59:698–717.

53. Sun Y, Xian L, Xing H, et al. Factors influencing the nuclear targeting ability of nuclear localization signals. J Drug Target. 2016;24:927–933.

54. Won YW, Lim KS, Kim YH. Intracellular organelle-targeted non-viral gene delivery systems. J Control Release. 2011;152:99–109.

55. Kim YH, Han ME, Oh SO. The molecular mechanism for nuclear transport and its application. Anat Cell Biol. 2017;50:77–85.

56. Aronsohn AI, Hughes JA. Nuclear localization signal peptides enhance cationic liposome-mediated gene therapy. J Drug Target. 1998;5:163–169.

57. Dominguez-Berrocal L, Cirri E, Zhang X, et al. New Therapeutic Approach for Targeting Hippo Signalling Pathway. Sci Rep. 2019;9:4771.

58. Hebert E. Improvement of exogenous DNA nuclear importation by nuclear localization signal-bearing vectors: a promising way for non-viral gene therapy? Biol Cell. 2003;95:59–68.

59. Lang A, Oye A, Eriksson J, Rowe AD, Lang E, Boe SO. Influence of acute promyelocytic leukemia therapeutic drugs on nuclear pore complex density and integrity. Biochem Biophys Res Commun. 2018;499:570–576.

60. Zhang F, White RL, Neufeld KL. Phosphorylation near nuclear localization signal regulates nuclear import of adenomatous polyposis coli protein. Proc Natl Acad Sci U S A. 2000;97:12577–12582.

61. Bolhassani A, Jafarzade BS, Mardani G. In vitro and in vivo delivery of therapeutic proteins using cell penetrating peptides. Peptides. 2017;87:50–63.

62. Efthymiadis A, Briggs LJ, Jans DA. The HIV-1 Tat nuclear localization sequence confers novel nuclear import properties. J Biol Chem. 1998;273:1623–1628.

63. Kristensen M, Birch D, Morck Nielsen H. Applications and Challenges for Use of Cell-Penetrating Peptides as Delivery Vectors for Peptide and Protein Cargos. Int J Mol Sci. 2016;17.

64. Hu M, Chen P, Wang J, Scollard DA, Vallis KA, Reilly RM. 123I-labeled HIV-1 tat peptide radioimmunoconjugates are imported into the nucleus of human breast cancer cells and functionally interact in vitro and in vivo with the cyclin-dependent kinase inhibitor, p21(WAF-1/Cip-1). Eur J Nucl Med Mol Imaging. 2007;34:368–377.

65. Niesner U, Halin C, Lozzi L, et al. Quantitation of the tumor-targeting properties of antibody fragments conjugated to cell-permeating HIV-1 TAT peptides. Bioconjug Chem. 2002;13:729–736.

66. Bauerle M, Doenecke D, Albig W. The requirement of H1 histones for a heterodimeric nuclear import receptor. J Biol Chem. 2002;277:32480–32489.

67. Jakel S, Mingot JM, Schwarzmaier P, Hartmann E, Gorlich D. Importins fulfil a dual function as nuclear import receptors and cytoplasmic chaperones for exposed basic domains. EMBO J. 2002;21:377–386.

68. Chuderland D, Konson A, Seger R. Identification and characterization of a general nuclear translocation signal in signaling proteins. Mol Cell. 2008;31:850–861.

69. Saijou E, Itoh T, Kim KW, Iemura S, Natsume T, Miyajima A. Nucleocytoplasmic shuttling of the zinc finger protein EZI Is mediated by importin-7-dependent nuclear import and CRM1-independent export mechanisms. J Biol Chem. 2007;282:32327–32337.

70. Beaudoin S, Paquette M, Fafard-Couture L, et al. Initial Evaluation of Antibody-conjugates Modified with Viral-derived Peptides for Increasing Cellular Accumulation and Improving Tumor Targeting. J Vis Exp. 2018.

71. Chauvin A, Boisvert FM. Proteomics Analysis of Colorectal Cancer Cells. Methods Mol Biol. 2018;1765:155–166.

72. Perez-Riverol Y, Csordas A, Bai J, et al. The PRIDE database and related tools and resources in 2019: improving support for quantification data. Nucleic Acids Res. 2019;47:D442–D450.

73. UniProt Consortium T. UniProt: the universal protein knowledgebase. Nucleic Acids Res. 2018;46:2699.

74. Choi H, Larsen B, Lin ZY, et al. SAINT: probabilistic scoring of affinity purification-mass spectrometry data. Nat Methods. 2011;8:70–73.

75. Montojo J, Zuberi K, Rodriguez H, et al. GeneMANIA Cytoscape plugin: fast gene function predictions on the desktop. Bioinformatics. 2010;26:2927–2928.

76. Thul PJ, Akesson L, Wiking M, et al. A subcellular map of the human proteome. Science. 2017;356.

77. Uhlen M, Fagerberg L, Hallstrom BM, et al. Proteomics. Tissue-based map of the human proteome. Science. 2015;347:1260419.

78. Uhlen M, Zhang C, Lee S, et al. A pathology atlas of the human cancer transcriptome. Science. 2017;357.

