## Supplemental figures for "Proteomic-based evaluation of nuclear transport of NLS-tagged trastuzumab-emtansine with enhanced cytotoxic potency"

### Slide 1
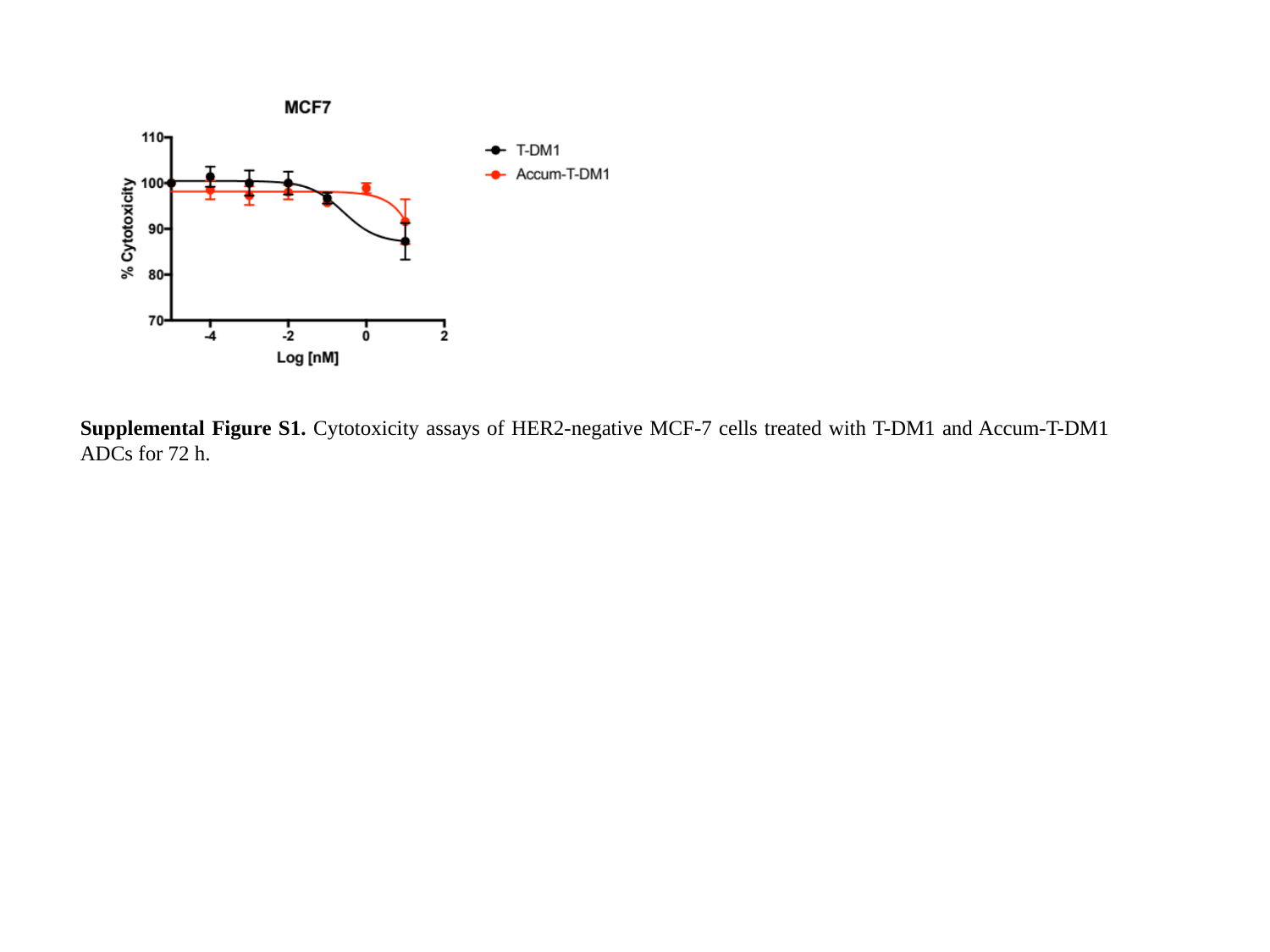

Supplemental Figure S1. Cytotoxicity assays of HER2-negative MCF-7 cells treated with T-DM1 and Accum-T-DM1 ADCs for 72 h.

### Slide 2
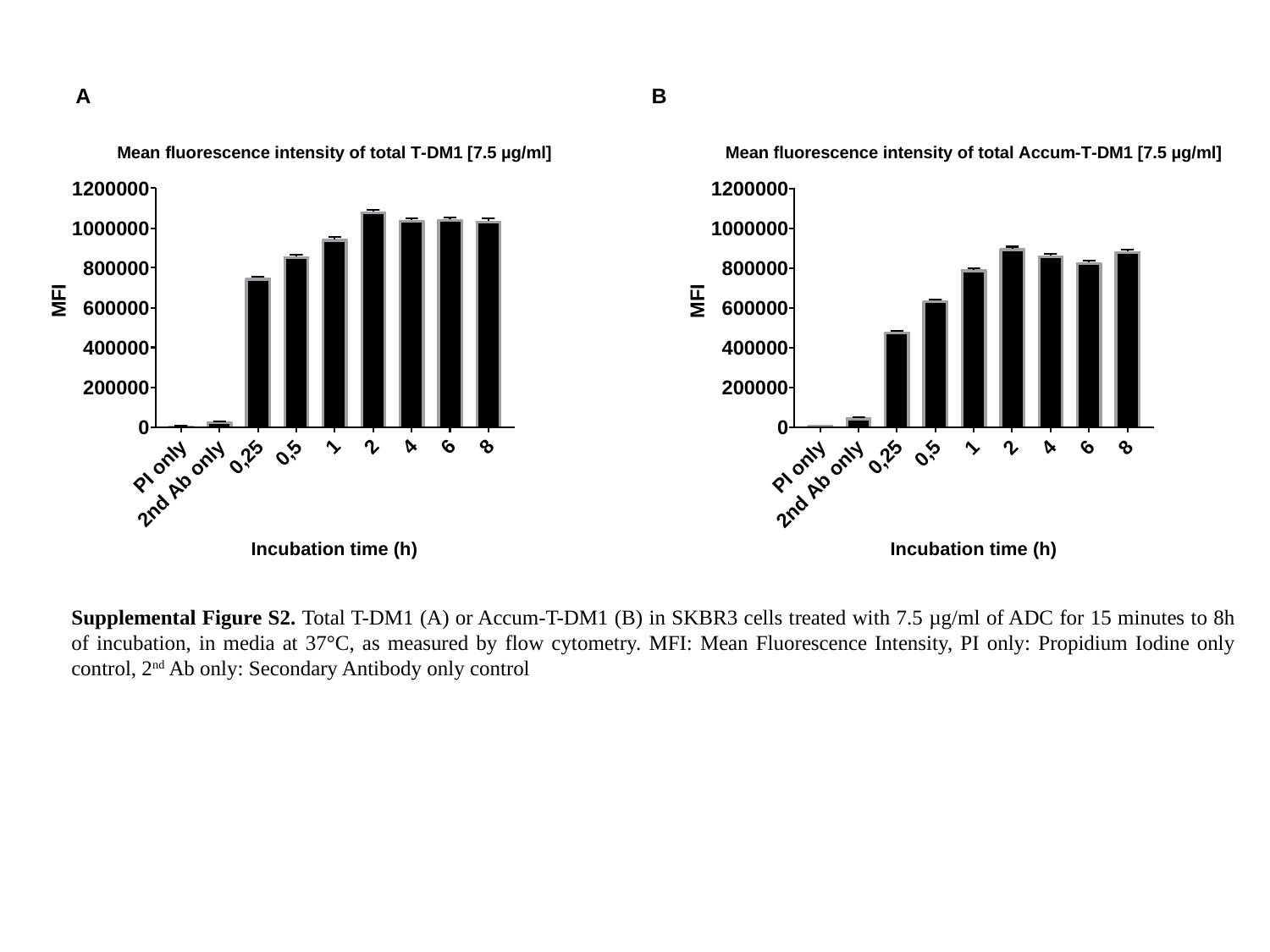

A
B
Supplemental Figure S2. Total T-DM1 (A) or Accum-T-DM1 (B) in SKBR3 cells treated with 7.5 µg/ml of ADC for 15 minutes to 8h of incubation, in media at 37°C, as measured by flow cytometry. MFI: Mean Fluorescence Intensity, PI only: Propidium Iodine only control, 2^nd^ Ab only: Secondary Antibody only control

### Slide 3
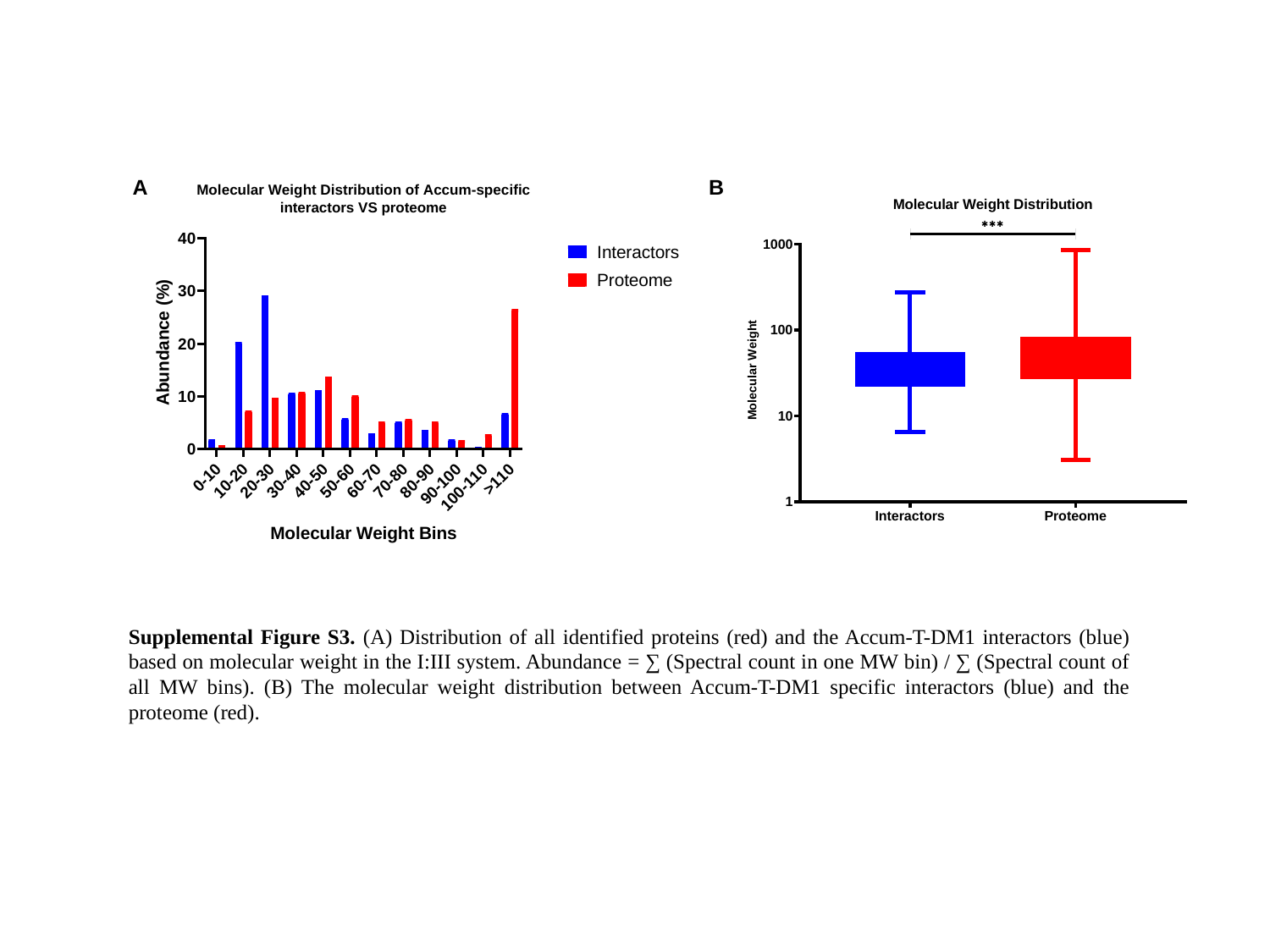

A
B
Supplemental Figure S3. (A) Distribution of all identified proteins (red) and the Accum-T-DM1 interactors (blue) based on molecular weight in the I:III system. Abundance = ∑ (Spectral count in one MW bin) / ∑ (Spectral count of all MW bins). (B) The molecular weight distribution between Accum-T-DM1 specific interactors (blue) and the proteome (red).

### Slide 4
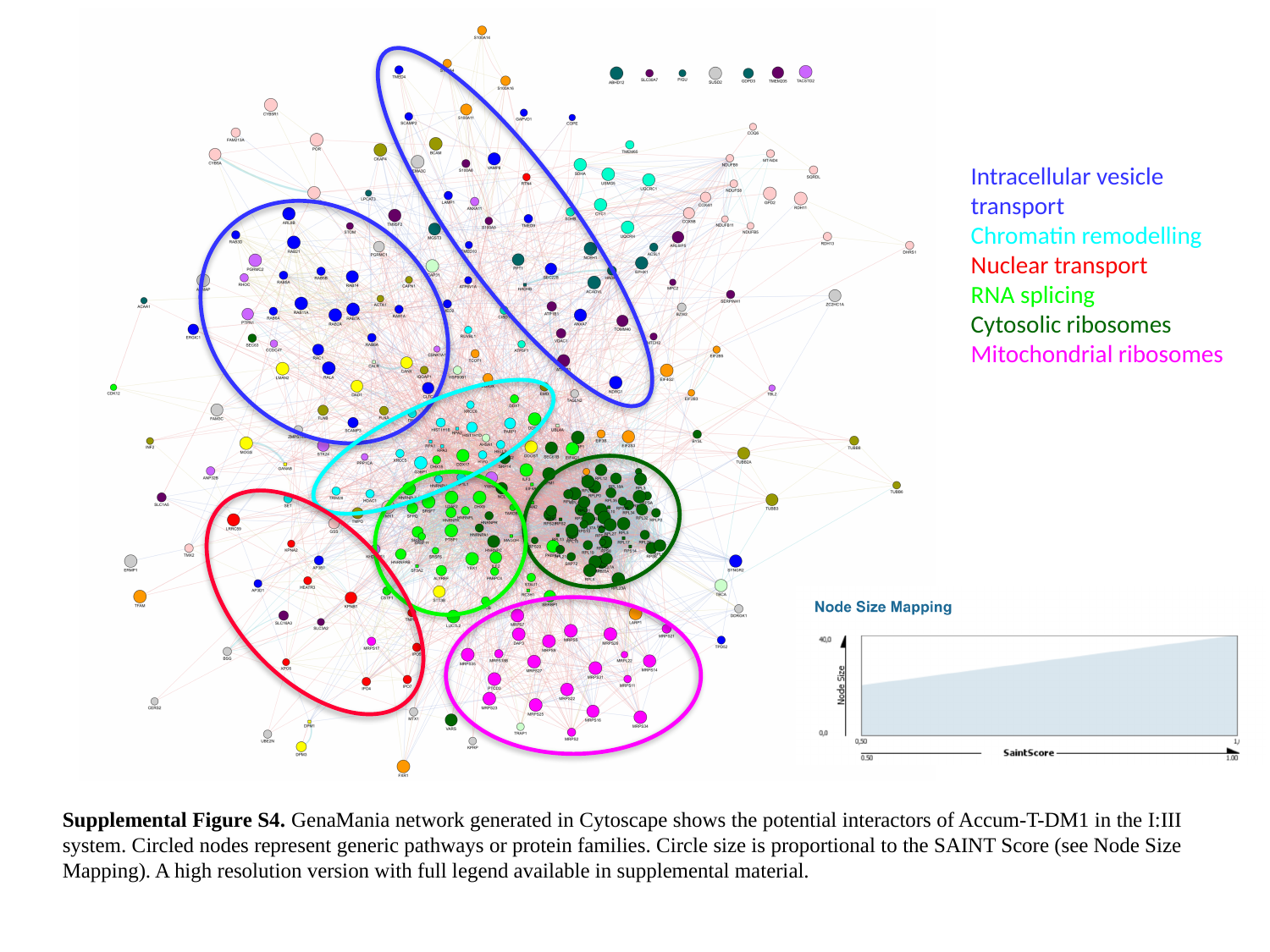

Intracellular vesicle transport
Chromatin remodelling
Nuclear transport
RNA splicing
Cytosolic ribosomes
Mitochondrial ribosomes
Supplemental Figure S4. GenaMania network generated in Cytoscape shows the potential interactors of Accum-T-DM1 in the I:III system. Circled nodes represent generic pathways or protein families. Circle size is proportional to the SAINT Score (see Node Size Mapping). A high resolution version with full legend available in supplemental material.

### Slide 5
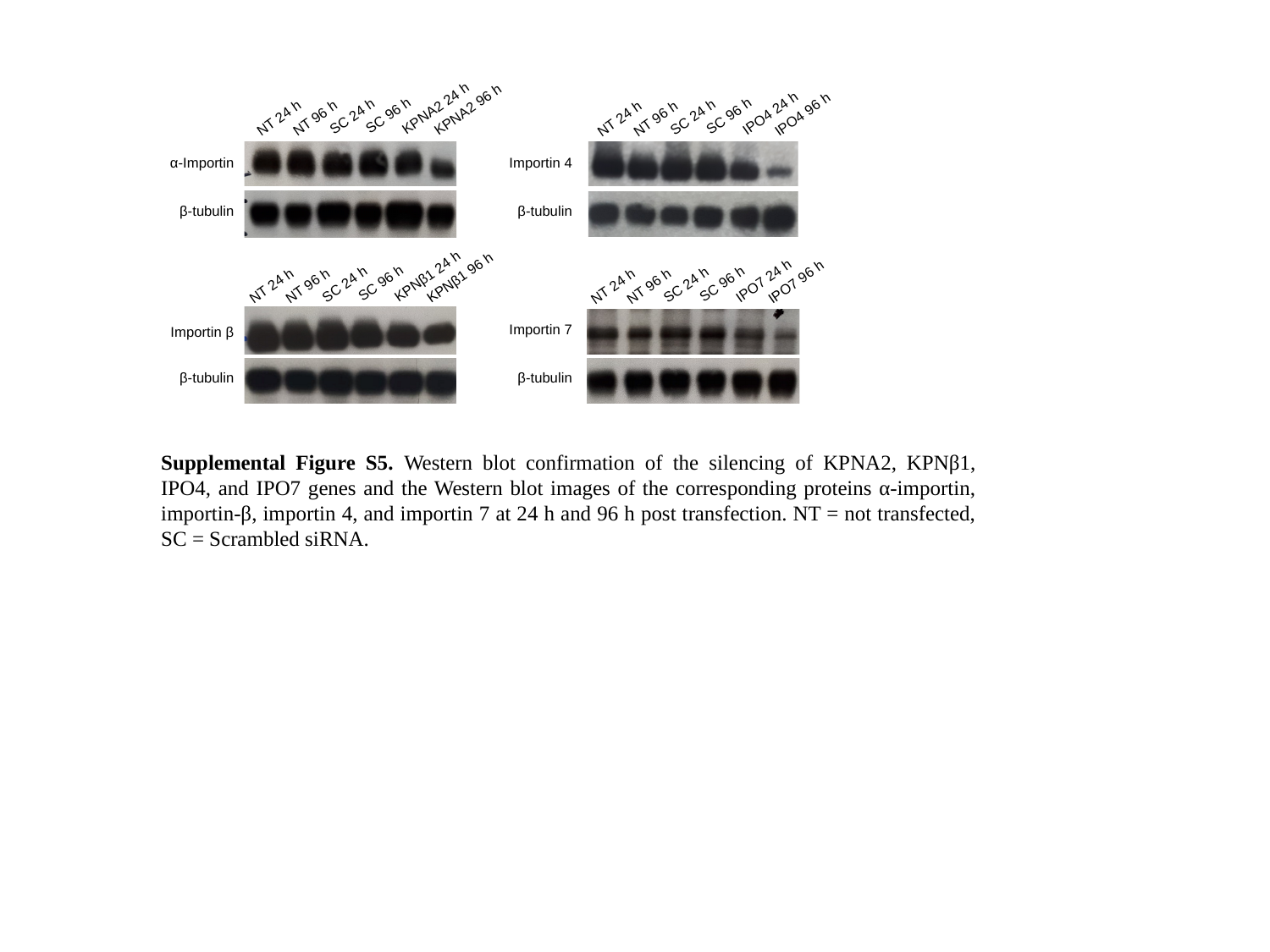

KPNA2 24 h
IPO4 24 h
KPNA2 96 h
IPO4 96 h
SC 96 h
SC 96 h
SC 24 h
SC 24 h
NT 24 h
NT 96 h
NT 24 h
NT 96 h
α-Importin
Importin 4
β-tubulin
β-tubulin
KPNβ1 24 h
IPO7 24 h
KPNβ1 96 h
IPO7 96 h
SC 96 h
SC 96 h
SC 24 h
SC 24 h
NT 24 h
NT 96 h
NT 24 h
NT 96 h
Importin 7
Importin β
β-tubulin
β-tubulin
Supplemental Figure S5. Western blot confirmation of the silencing of KPNA2, KPNβ1, IPO4, and IPO7 genes and the Western blot images of the corresponding proteins α-importin, importin-β, importin 4, and importin 7 at 24 h and 96 h post transfection. NT = not transfected, SC = Scrambled siRNA.
